# Structural Basis of Mitochondrial Receptor Binding and GTP-Driven Conformational Constriction by Dynamin-Related Protein 1

**DOI:** 10.1101/172809

**Authors:** Raghav Kalia, Ray Yu-Ruei Wang, Ali Yusuf, Paul V. Thomas, David A. Agard, Janet M. Shaw, Adam Frost

## Abstract

Mitochondrial inheritance, genome maintenance, and metabolic adaptation all depend on organelle fission by <u>D</u>ynamin-<u>R</u>elated <u>P</u>rotein 1 (DRP1) and its mitochondrial receptors. DRP1 receptors include the paralogs <u>M</u>itochondrial <u>D</u>ynamics 49 and 51 (MID49/MID51) and <u>M</u>itochondrial <u>F</u>ission <u>F</u>actor (MFF), but the mechanisms by which these proteins recruit DRP1 and regulate its activities are unknown. Here we present a cryoEM structure of human, full-length DRP1 bound to MID49 and an analysis of structure- and disease-based mutations. We report that GTP binding allosterically induces a remarkable elongation and rotation of the G-domain, Bundle-Signaling Element (BSE) and connecting hinge loops of DRP1. In this nucleotide-bound conformation, a distributed network of multivalent interactions promotes DRP1 copolymerization into a linear filament with MID49, MID51 or both. Subsequent GTP hydrolysis and exchange within the filament leads to receptor dissociation, shortening through disassembly, and concomitant curling of DRP1 oligomers into closed rings. The dimensions of the closed DRP1 rings are consistent with DRP1-constricted mitochondrial tubules observed in human cells. These structures are the first views of full-length, receptor- and nucleotide-bound dynamin-family GTPases and—in comparison with nucleotide-free crystal structures—teach us how these molecular machines perform mechanical work through nucleotide-driven allostery.

## Introduction

Pioneering work in yeast and other model systems revealed that fragmentation of the mitochondrial reticulum disperses units of the organelle during cell division^1–3^, coordinates morphological adaptation with metabolic demand^4–7^, and quarantines damaged units for turnover^8–10^. Recent work also led to the discovery of the role mitochondrial fission plays in regulated cell death pathways^11–19^, brain development and synaptic function^20–23^, and how certain pathogens disrupt these processes and hijack mitochondrial resources^24–26^. Finally, there is a growing understanding of how interorganelle contacts between the ER and mitochondria initiate mitochondrial fission^27,28^, and how this process regulates mitochondrial genome duplication and integrity^29,30^. The master regulator that unites these processes across eukaryotic evolution is the membrane-remodeling GTPase DRP1^4,7,31–37^.

DRP1 is necessary but not sufficient for mitochondrial fission because receptor proteins must recruit the enzyme to the <u>O</u>uter <u>M</u>itochondrial <u>M</u>embrane (OMM). In mammals, these include the paralogs <u>MI</u>tochondrial <u>D</u>ynamics proteins MID49 and MID51 or the <u>M</u>itochondrial <u>F</u>ission <u>F</u>actor, MFF ^32,35,38–41^. Following receptor-dependent recruitment, DRP1 assembles into polymers that encircle mitochondria and, through still poorly understood mechanisms, channels energy from GTP binding, hydrolysis, and nucleotide exchange into a mechanochemical constriction^13,41–46^. In addition to DRP1 and its OMM receptors, a recent study revealed that a second member of the dynamin-family of GTPases, dynamin-2, enacts the final fission event downstream of DRPI-driven constriction of a mitochondrial tubule^47^. Thus, mitochondrial division is a stepwise reaction regulated by DRP1 receptor binding, oligomerization and nucleotide-dependent conformational dynamics.

We and others have reported that the OMM receptors MFF or MID49/51 are independently sufficient to recruit DRP1 in order to divide mitochondria^38,39,41,48^. We also reported that MID49/51 can coassemble with DRP1 forming a copolymer with a dramatically different morphology than reported for dynamin-family members^41^. While these results suggest that an adaptor protein could alter the architecture of a dynamin polymer to facilitate a mitochondria-specific activity, the architecture and functions of this coassembly remain unclear.

Here, we report the structural basis of DRP1 coassembly with MID49/MID51. Our cryoEM structure reveals how nucleotide binding to the G-domain induces conformational changes that allosterically propagate through the BSE to open and elongate DRP1 and thereby expose multiple receptor-binding surfaces. MID49/51 binding stabilizes a specific alignment of DRP1 tetramers to nucleate polymerization of a linear co-filament. Then, in a path-dependent reaction, we show how GTP hydrolysis and nucleotide exchange lead to conformational constriction by the polymer. Specifically, when DRP1 subunits within the cofilament exchange and hydrolyze nucleotide, they dissociate from MID49/51 receptors and the entire polymer dynamically shortens and curls into a closed ring. Analysis of structure-based and disease-causing mutations indicate that allosterically driven rearrangements of the stalk helices—and the critical L1N^S^ loop—support curling from linear strings into constricted rings following receptor dissociation.

## Results and Discussion

To date, many structural studies of dynamin-family proteins have relied upon mutated or truncated constructs to facilitate crystallization. We purified wild-type, full-length human DRP1 including the N-terminal GTPase domain (G-domain), Bundle Signaling Element (BSE), and four-helix bundle known as the stalk (Fig. 1a). This construct also contained the lipid binding ~100 amino acid region referred to as the variable domain (VD) that sits between the third and fourth alpha helices of the stalk, analogous to the Pleckstrin Homology (PH) domain found in endocytic dynamin proteins. A crystal structure of a nucleotide-free and truncated DRP1 mutant revealed the organization of these domains and an overall similarity with the structure of nucleotide-free endocytic dynamin^42,49,50^. We also purified soluble truncations of MID49 and MID51 engineered to lack their N-terminal transmembrane anchors but include the cytoplasmic nucleotidyltransferase-like domain and the “<u>D</u>ynamin <u>R</u>ecruitment <u>R</u>egion” (DRR) required for DRP1 binding (Fig 1a)^38,51–54^.

**Figure 1:**
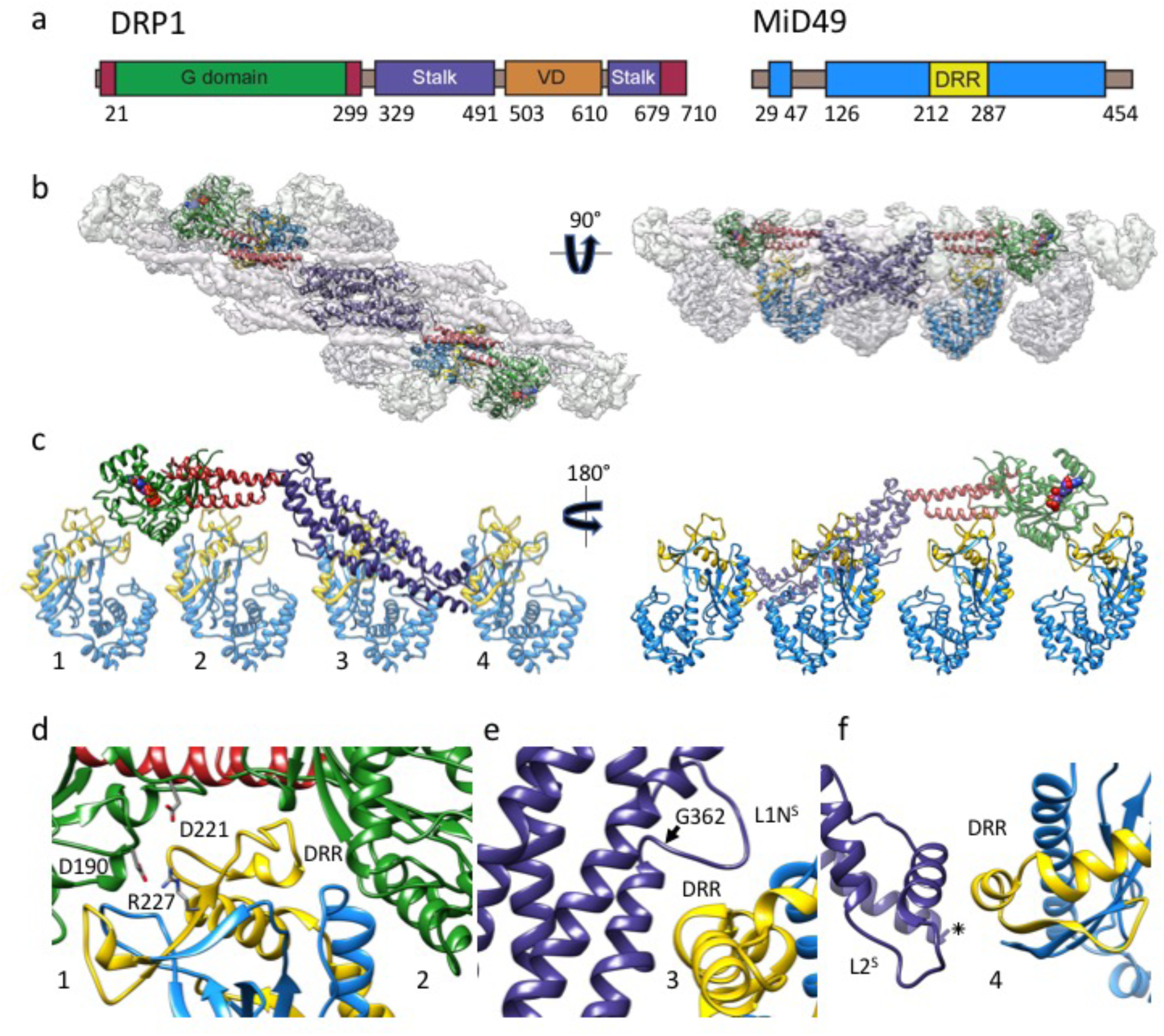
Architecture of the DRP1-MID49 linear filament: (**a**) DRP1 and MID49 domain arrangements. (**b**) Density map and atomic models for DRP1 and MID49_126-454_. Green: G-domain, Red: Bundle Signaling Element (BSE), Purple: Stalk, Blue: MID49, Yellow: Dynamin Recruitment Region (DRR) of MID49. (**c**) Each DRP1 chain contacts four different MID49 molecules, as numbered. (**d**) MID-interaction surfaces 1 and 2. Green ribbons on either side of the DRR come from two separate G-domains, and the residues involved in a key salt bridge for MID-interaction surface 1 are shown as sticks. (**e**) Rotated view of MID-interaction surface 3. Disease-associated DRP1 residue G362 (arrow) supports the conformation of the L1N^S^ loop essential for linear copolymerization with MID receptors. (**f**) Rotated view of MID-interaction surface 4 and the L2^S^ loop.

Incubating equimolar ratios of DRP1 with soluble MID49_126-454_, MID51_132-463_, or both proteins together, in the presence of GTP resulted in cofilament assembly (Extended Data Figs. 1–2). We focused on the filaments formed with MID49_126-454_ in the presence of the non-hydrolyzable analog GMPPCP and determined their structure from cryoEM images (Extended Data Figs. 3–9 and Table 1). 3D reconstruction resolved elongated DRP1 dimers bound stoichiometrically to MID49_126-454_, without assignable density for the variable domain (Fig. 1b, Extended Data Fig. 4). Surprisingly, each chain of DRP1 bound MID49 at four different sites, and each MID49 in turn bound four DRP1 molecules to yield a vast and highly avid interaction network (Figs. 1b-c, Extended Data Figs. 4 and 8). MID49 binding to four separate DRP1 molecules stabilized a linear arrangement of inter-DRP1 interfaces reminiscent of those observed for other dynamin-family proteins by X-ray crystallography (Fig. 1b, Extended Data Figs. 4 and 8)^42,49,50,55–57^.

MID49’s DRR motif occupied the space between two neighboring G-domains and contacted both via MID-interaction interfaces 1 and 2 (buried surface areas of ~530Å^2^ and ~200 Å^2^, respectively, see Fig. 1b-d). The precise spacing required for this bivalent G-domain interaction explains why previous mutagenesis efforts suggested that the size and topology of the β4–α4 loop, rather than its exact sequence (which differs between MID49 and MID51) are critical determinants of binding. Single point mutations within this loop for MID49 and MID51 did not disrupt binding, while mutations that altered the length, topology or positioning of the loop did^51,52,54^. We found that mutating conserved DRP1 residues involved in interface 1—the energetically most significant interface—did prevent coassembly. Specifically, the D190A mutation should neutralize an interface 1 salt bridge and the D221A mutation should alter the conformation of an interface 1 loop (Fig. 1d). We observed that both mutants prevented assembly with MID49 and altered DRP1’s self-assembly properties (Extended Data Figs. 10–12).

Unexpectedly, MID49 also made contact with the stalk loops of a third and fourth DRP1 molecule through MID-interaction interfaces 3 and 4 (buried surface areas of ~450Å^2^ and ~230Å^2^, respectively, Fig. 1c, e, and f). The loops involved in both interaction interfaces 3 and 4 determine the assembly properties of higher-order dynamin-family oligomers ^42,49,50,55–57^. MID-interaction surface 3, in particular, harbors the conserved loop L1N^S^ and is the site of multiple disease alleles that lead to elongated mitochondrial morphology, including G362D and G363D (Fig. 1e, Extended Data Figs. 10–11 and ^58–60^). Prior work has also established that this loop comprises part of the intra-molecular PH domain binding site for the soluble state of endocytic dynamin tetramers (Extended Data Fig. 13)^57^, and is a determinant of conformational heterogeneity for these and other dynamin-family proteins^55,57,61^. The presence of disease alleles near this interface suggests that these mutations may compromise receptor interactions and that defects in the recruitment of DRP1 to mitochondria may contribute to pathogenesis. As discussed below, we found that the G362D mutant (Fig. 1e) failed to coassemble with MID49 and displayed altered assembly and conformational properties (Extended Data Figs. 12–15).

Understanding the allosteric coupling between nucleotide binding, hydrolysis or exchange and the conformational repertoire of dynamin-family GTPases remains an unmet challenge^62,63^. We observed that the GMPPCP-bound G-domains and the BSE of DRP1 adopt strikingly different conformations in the cryoEM density compared to the nucleotide-free crystal structure^42^. In addition to other nucleotide-induced conformational changes within the G-domain, the most salient are the closing of the G2/switch-1 loop to form a closed “lid” over the nucleotide (Figs. 2a-c). The closure of the switch-1 lid propagates through the adjacent beta sheet to push the α-helices of the BSE into an orthogonal position (Extended Data Movie 1). When evaluated in the context of a stalk interface-2 DRP1 dimer, this conformational change is an impressive 90° rotation of the G-domain and a 40Å translation toward the stalk (Fig. 2d, Extended Data Movies 2-3). Two of the three DRP1 surfaces that engage the DRR of MID49 (MID receptor interaction interfaces 1 and 2, Figs. 1–2) are inaccessible in the nucleotide-free state but become available for binding upon nucleotide-driven elongation (Fig. 2d, Extended Data Movies 2-3).

**Figure 2:**
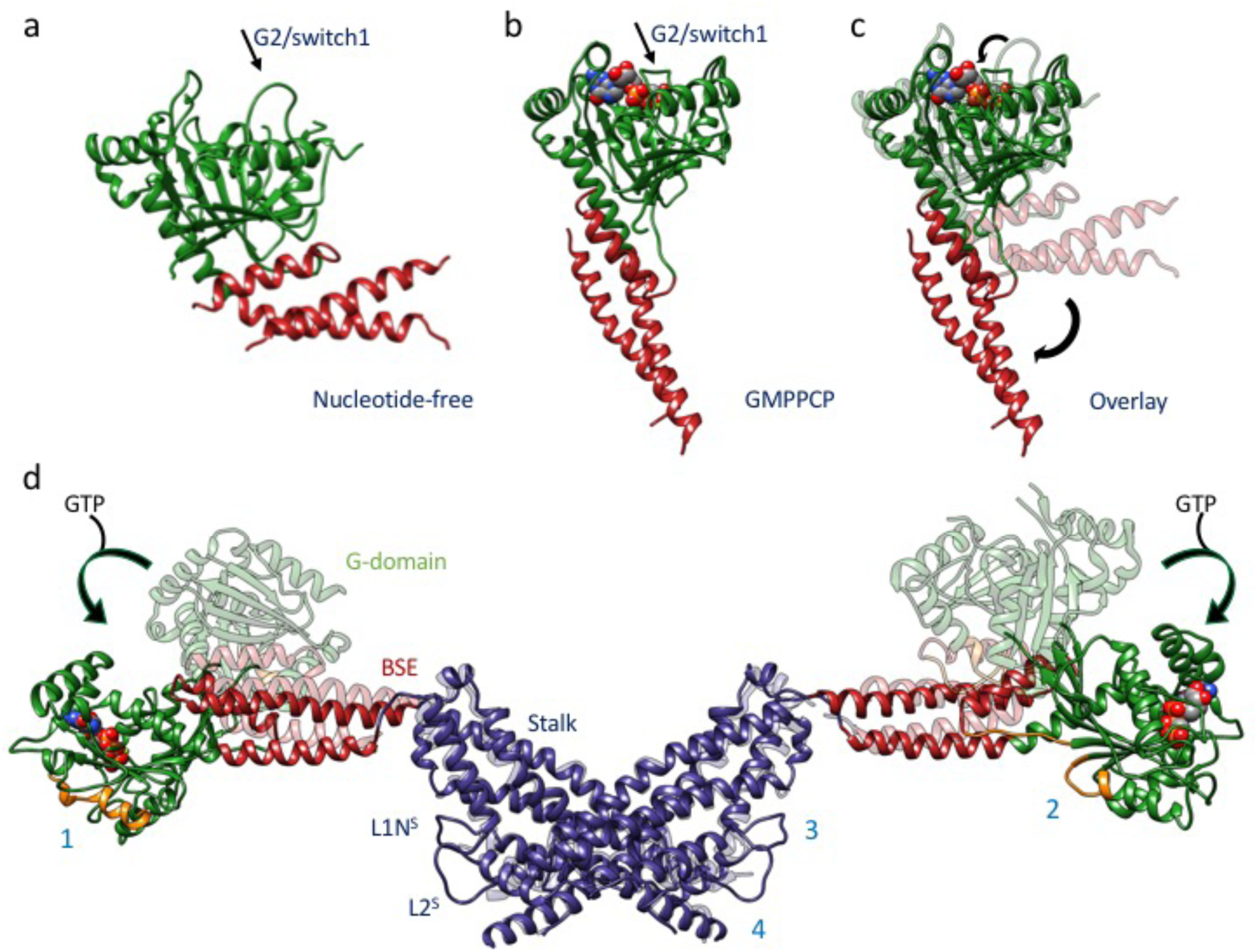
Nucleotide-driven allosteric elongation of DRP1 exposes MID binding sites: **(a)** nucleotide-free state of DRP1 G-domain and BSE as seen in a crystal structure (PDB ID:4BEJ). Arrow points to the G2/switch1 loop. **(b)** GMPPCP-bound G-domain-BSE conformation determined by cryoEM. **(c)** Overlay of A and B. Curved arrows highlight the closing of G2/switch 1 “lid” and the opening of the BSE “wrist”. **(d)** Global conformational change induced by nucleotide binding. Rotation and translation of the G-domain and BSE elongates the dimer and exposes MID-interaction surfaces 1 and 2 (annotated on separate monomers for clarity). The surfaces of the G-domains that engage MID receptors are rendered orange in the nucleotide-bound and elongated conformation.

Since DRP1’s ability to polymerize into linear filaments with MID49 or MID51 depended on non-hydrolyzable GTP analogs, we next evaluated the dynamics of these filaments in the presence of hydrolyzable GTP. Following copolymerization in the presence of the non-hydrolyzable analog, we exchanged GMPPCP for GTP through dialysis and followed the reaction using negative stain transmission electron microscopy at sequential time points until the GTP was exhausted. We observed that the linear DRP1-MID49 cofilaments disassembled into small oligomers upon complete hydrolysis to GDP and exhibited a fascinating dynamic instability at intermediate time points (Fig. 3). Specifically, the long and linear filaments seen at early time points disassembled into shorter, curling oligomers that—upon reaching a reproducibly narrow range of lengths—spontaneously closed into constricted rings that were remarkably uniform in diameter (Fig. 3).

**Figure 3:**
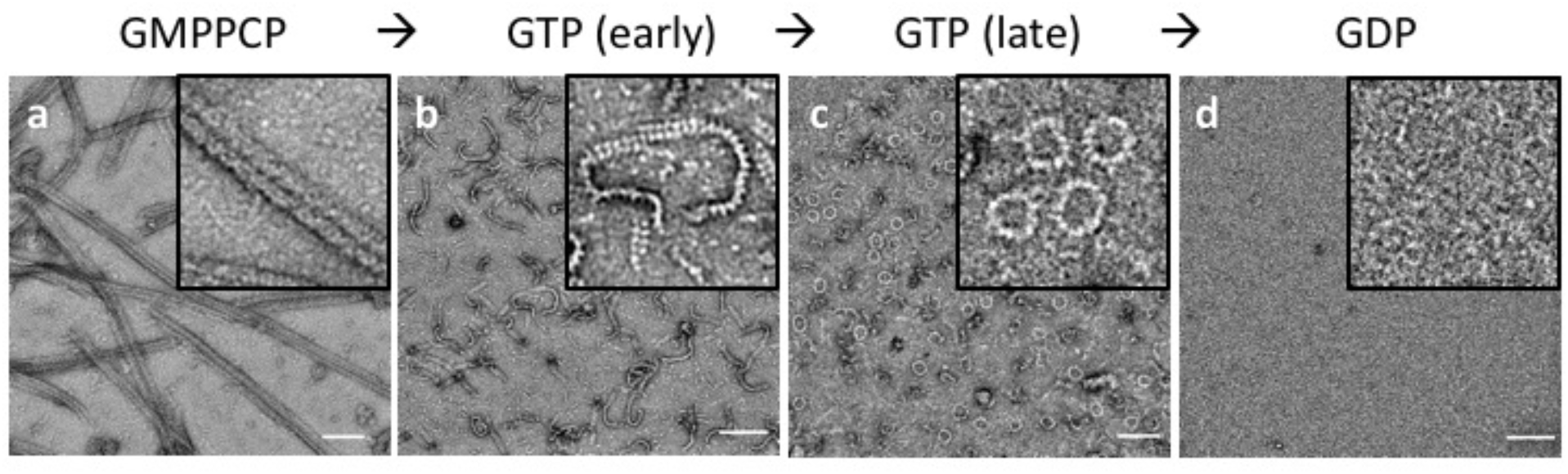
Dynamic instability of the DRP1-MID49 linear assembly and curling into closed DRP1 rings: **(a)** DRP1-MID49_126-454_ linear filaments copolymerized with GMPPCP. **(b-c)** Subsequent exchange into GTP leads to partial disassembly and curling into closed rings. **(d)** GTP exhaustion leads to complete oligomer disassembly. Bars=100nm.

In a separate but related experiment, we evaluated the assembly properties of the DRP1 mutant G362D with and without MID49_126-454_. As described above, this disease-associated residue sits at the base of the L1N^S^ loop that forms part of the third interface with MID49 (Figs. 1c, 1e, Extended Data Fig. 13). We found that DRP1_G362D_ purified as a nearly monodisperse and stable dimer, rather than a mix of tetramers and higher order species observed for the wild-type, full-length protein (Extended Data Fig. 12). In addition, DRP1_G362D_ exclusively formed rings, not filaments, with or without MID49_126-454_ and in the presence of GMPPCP or GTP (Fig. 4a, Extended Data Figs. 14–15). These rings resembled those observed with wild-type DRP1 in all respects except that the wild-type protein only formed closed rings from the pre-formed linear MID49 copolymers through the path-dependent reaction described above (Fig. 3 versus Fig. 4). Using the non-hydrolyzable GTP analog GMPPCP with the DRP1_G362D_ rings also improved structural homogeneity, presumably because the wild-type rings remain dynamic and eventually disassembled in the presence of GTP upon complete hydrolysis to GDP (Fig. 3).

**Figure 4:**
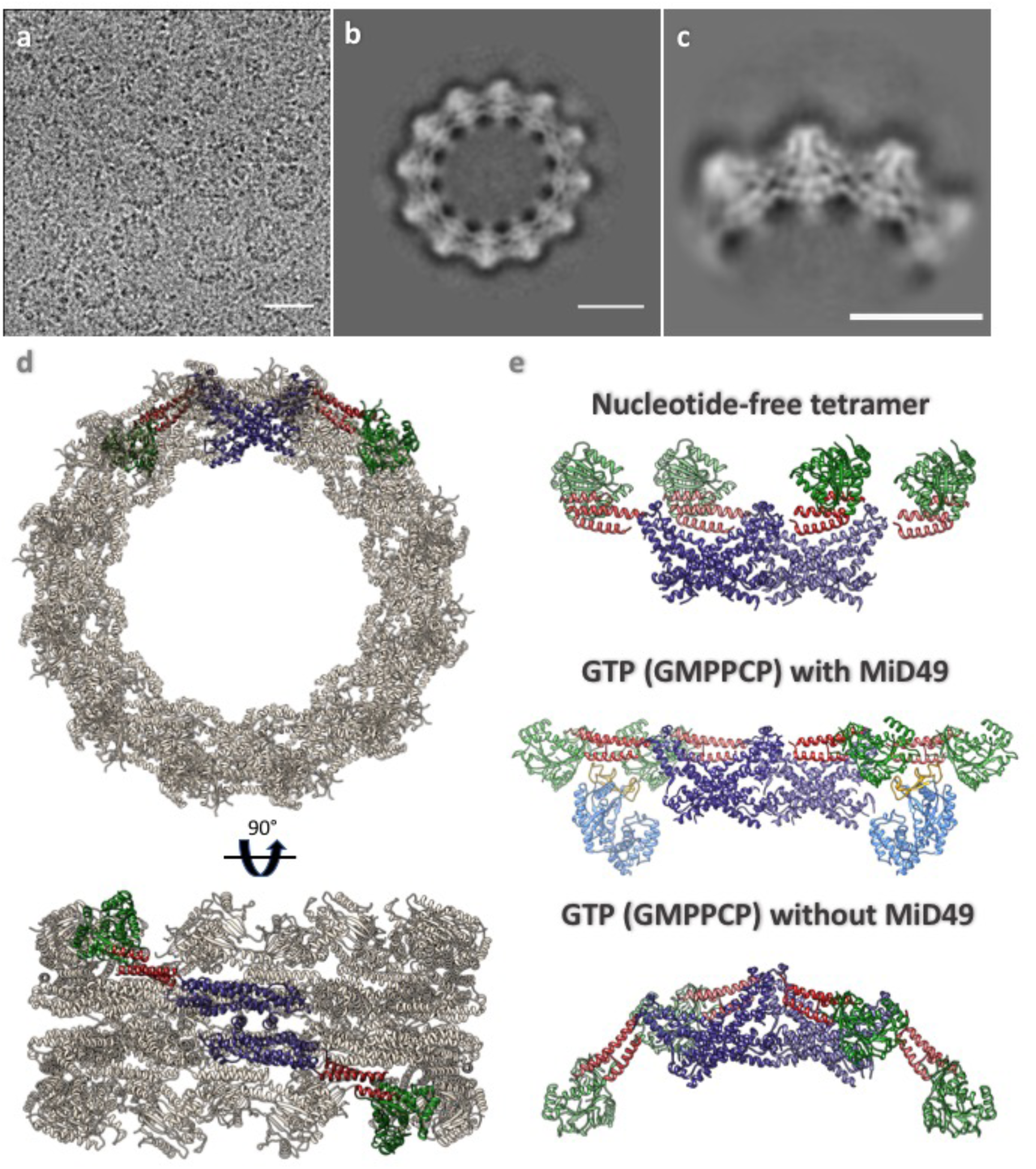
DRP1_G362D_ cannot bind MID49 and forms rings exclusively with GMPPCP or GTP. **(a)** CryoEM micrograph of DRP1_G362D_ rings. **(b)** 2D class average of the predominant closed ring that comprises 12 DRP1 dimers. **(c)** 2D class average of a quarter of the ring revealing the secondary structure elements of the “X”-shaped DRP1 dimer. **(d)** 3D model of the closed ring. **(e)** Comparison between DRP1 tetramers observed in the nucleotide-free state (top, PDB ID:4BEJ), the GMPPCP and MID49_126-454_-bound linear state (middle), and the bent conformation modeled for the rings. Bar for **(a)** = 30nm, for **(b, c)** =100Å.

We imaged the DRP1_G362D_ rings using cryoEM and used 2D class averages of the predominant 12-dimer closed ring to model the architectural differences between the linear filaments and the closed rings (Fig. 4, Extended Data Fig. 14–15). To account for the projected ring density, the G-domain and the BSE of DRP1 must move even further down toward the stalks. Moreover, while stalk interface-2 remains constant—as revealed by the X-shaped projected dimer (Fig. 4c)—the curvature of the ring dictates that stalk interfaces 1 and 3 must be extensively remodeled to allow a ~30 degree bending per dimer in comparison with the linear DRP1-MID49 copolymer (Fig. 4d, Extended Data Fig. 15, Movie 4). We did not observe any density for MID49 in the wild-type rings that form by curling of the MID49-DRP1 cofilament in the presence of GTP, nor in our higher-resolution analysis of the DRP1_G362D_ rings that form with or without MID49 present (Fig. 4b-c, Extended Data Fig. 14–15).

We note that even the most constricted form of the closed ring is insufficient to drive complete mitochondrial fission because the inner diameter is only ~16nm. This length suggests that a constricted membrane tubule would be stable and that the structures we have observed in vitro could correspond with the highly-constricted but pre-fission state observed in living cells when another dynamin-family protein, the primarily endocytic dynamin-2, is depleted^47^. Thus, initial constriction by DRP1 may stabilize the high degree of membrane curvature that is suitable, perhaps even tuned, for the recruitment and final fission event catalyzed by additional dynamin-family enzymes.

Together, these findings establish four conceptual advances. First, our cryoEM structure revealed how receptor proteins like MID49/MID51 recruit and stabilize a specific nucleotide-bound conformation of DRP1 and nucleate polymerization of a cofilament. We speculate that the nearly linear properties of this polymer have adapted to encircle low-curvature mitochondrial tubules. The selective stabilization of this open, elongated conformation of DRP1 within the MID receptor cofilament explains why overexpression of the MID receptors inhibits mitochondrial fission^39^. Second, analysis of the MID49-DRP1 copolymer exposed how nucleotide binding induces an impressive conformational rearrangement to expose a network of multi-valent receptor binding sites. We now understand these nucleotide-driven allosteric transformations in detail and in the context of full-length and oligomeric DRP1. Third, a path-dependent constriction reaction revealed intrinsic GTP-dependent DRP1 properties that are reminiscent of microtubule dynamic instability^64^. In this reaction, nucleotide exchange and hydrolysis led to MID49/51 receptor dissociation, disassembly from the ends of the linear filament, and concomitant curling of the shortened filaments into closed rings. Fourth and finally, analysis of a disease mutant in the L1N^S^ loop, DRP1_G362D_, exposed this loop as a vital determinant of mitochondrial receptor binding and the dynamic inter-stalk interactions that govern oligomer architecture and the ability of dynamin proteins to perform mechanical work.

## Acknowledgments

We thank Michael Braunfeld, Cameron Kennedy, David Bulkley, and Alexander Myasnikov and the UCSF Center for Advanced CryoEM, which is supported in part from NIH grants S10OD020054 and 1S10OD021741 and the Howard Hughes Medical Institute. We also thank the QB3 shared cluster and NIH grant 1S10OD021596-01, Jean-Paul Armache, Nathaniel Talledge for microscopy advice, and Charles Greenberg for consulting on structural modeling. This work was further supported by a Faculty Scholar grant from the Howard Hughes Medical Institute (A.F.), the Searle Scholars Program (A.F.), NIH grant 1DP2GM110772-01 (A.F.), NIH grants GM53466 and GM84970 (J.S.), and the Howard Hughes Medical Institute (R.W., J.S., D.A.). A.F. is a Chan Zuckerberg Biohub investigator.

## Author Contributions

R.K., J.S. and A.F. conceived of the study. R.K., R.W., A.Y. and P.T. performed all experiments. R.K., R.W. and A.F. performed the computational analyses. All authors evaluated the results and edited the manuscript. R.K. and A.F. wrote the manuscript with input from all of the authors.

## Data Accessibility

All of the cryoEM density maps associated with this study have be deposited in the EMDB with accession numbers EMD-8874(PROC). The atomic coordinates have been deposited in the PDB as 5WP9(PROC).

## Extended Data

### Methods

#### Construct design

Wild type (WT) DRP1 isoform 2 sequence was purchased from DNASU (sequence ID HsCD00043627, UNIPROT identifier: O00429-3 also known as DLP1a) and was cloned into pET16b plasmid (Novagen) between the Nde1 and BamH1 sites. The vector was kindly provided by the laboratory of Wesley I. Sundquist with a 10X-His tag followed by a PreScission protease site (Leu-Glu-Val-Leu-Phe-Gln-Gly-Pro). Wild-type MID49_126-454_ sequence was PCR amplified and cloned into pGEX6p1 vector having an N-terminal GST tag followed by a PreScission protease site. Site directed mutagenesis was performed on pET16b-DRP1 using the Gibson cloning method to introduce mutations^1^. All constructs were verified using Sanger sequencing.

#### Protein purification

Protein purification was performed as described^2^. Briefly, plasmids containing the WT DRP1 or MID49_126-454_ sequence were transformed in the BL21-DE3 (RIPL) strain of *E. coli.* The colonies were inoculated in LB culture medium and grown overnight. Secondary inoculations were done the next morning in ZY medium for auto-induction^3,4^. The cultures were grown to an OD_600_ of 0.8 at 37°C in baffled flasks and were shifted to 19°C to grow for another 12 hours. The cultures were spun down and the bacterial pellets were used for protein purification immediately or stored at −80°C.

Full length DRP1 WT and mutant variants were purified as described previously for DRP1 WT with modifications^5^. Briefly, the bacterial pellets were resuspended in buffer A (50 mM HEPES/NaOH (pH 7.5), 400 mM NaCl, 5 mM MgCl_2_, 40 mM imidazole, 1 mM DTT, 0.5 mg DNase (Roche) and protease inhibitors (10 mM pepstatin, 50 mM PMSF, 0.5 mM aprotinin and 2 mM leupeptin), followed by cell disruption with a probe sonicator. Lysates were cleared by centrifugation at 40,000xg in Beckman JA 25.5 rotors for 60 min at 4°C. The supernatant was filtered using a 0.45 μm filter and applied to Ni-NTA Agarose beads pre-equilibrated with buffer B (50 mM HEPES/NaOH (pH 7.5), 400 mM NaCl, 5 mM MgCl_2_, 40 mM imidazole, 1 mM DTT). The beads were washed with 20 column volumes each of buffer B and buffer C (50 mM HEPES/NaOH (pH 7.5), 800 mM NaCl, 5 mM MgCfe, 40 mM imidazole, 1 mM DTT, 1 mM ATP, 10 mM KCl) followed by buffer D (50 mM HEPES/NaOH (pH 7.5), 400 mM NaCl, 5 mM MgCl_2_, 80 mM imidazole, 1 mM DTT, 0.5% (w/v) CHAPS). A final pre-elution wash was done with 20 column volumes of buffer B. Bound DRP1 was eluted with buffer E (50 mM HEPES/NaOH (pH 7.5), 400 mM NaCl, 5 mM MgCl_2_, 300 mM imidazole, 1 mM DTT) and dialyzed overnight at 4°C against buffer B without imidazole in the presence of PreScission protease to cleave the N-terminal 10X-His tag. The protein was re-applied to a Ni-NTA column pre-equilibrated with dialysis buffer and was observed to bind the column without the 10X-His tag as well. Subsequently, the protein was eluted with buffer B containing 80 mM imidazole.

Pure protein was concentrated with a 30kDa molecular weight cutoff (MWCO) centrifugal concentration device (Millipore). In a final step, DRP1 was purified by size-exclusion chromatography (SEC) on a Superdex-200 column (GE) in buffer F containing 20 mM HEPES/ NaOH (pH 7.5), 300 mM NaCl, 2.5 mM MgCl_2_ and 1 mM DTT. Fractions containing DRP1 were pooled, concentrated, flash frozen as single use aliquots in liquid nitrogen and stored at −80°C. Exact masses for purified DRP1 proteins were validated by MALDI-TOF mass spectrometry.

MID49_126-454_ was purified as described with modifications^6^. pGEX6p1-MID49_126-454_ plasmid DNA (human, UNIPROT identifier: Isoform 1 Q96C03-1, also known as MIEF2) was transformed in BL21 (DE3) RIPL cells. The colonies were grown overnight in LB medium and secondary cultures were grown in ZY medium. Cells were grown to an OD_600_ of 0.8-1, collected by centrifugation and processed immediately or stored at −80°C as described above. The bacterial pellets were lysed as described above in MID-buffer A (50 mM Tris pH 8.0, 500 mM NaCl, 5% glycerol, 1 mM DTT and 0.1% (v/v) Triton X-100). The lysates were pre-cleared at 40,000xg and filtered using a 0.45 μm filter before applying to 3 ml glutathione sepharose beads (GE). After overnight binding to beads, the unbound protein was removed and the beads were washed using 20 column volumes each of MID-buffer A and MID-buffer B (50 mM Tris pH 8.0, 500 mM NaCl, 5% glycerol, 1 mM DTT). The protein was eluted with MID-buffer C (50 mM Tris pH 8.0, 500 mM NaCl, 5% glycerol, 1 mM DTT and 20 mM reduced glutathione). The eluate was cleaved overnight with PreScission protease while dialyzing against MID-buffer D (20 mM Tris pH 8.0, 100 mM NaCl, 5% glycerol, 1 mM DTT). Cleaved protein was further purified using ion-exchange chromatography using a Q sepharose (GE) column. The low salt buffer for ion-exchange was the same as MID-buffer D and the high salt buffer was MID-buffer E (20 mM Tris pH 8.0, 1 M NaCl, 5% glycerol, 1 mM DTT). The relevant MID49_126-454_ fractions were pooled, concentrated and further purified using an SEC column pre-equilibrated with MID-buffer F (20 mM Tris pH 8.0, 200 mM NaCl, 5% glycerol, 1 mM DTT). The fractions containing MID49126-454 were pooled, concentrated, flash frozen in liquid nitrogen and stored as single use aliquots at −80°C.

#### Filament assembly, EM sample preparation, data acquisition and processing

To assemble DRP1-MID49_126-454_ filaments, the proteins were mixed to a final concentration of 2 μM each and kept for an hour at room temperature. The mixture was dialyzed against assembly buffer (20 mM HEPES pH 7.5, 50 mM KCl, 3 mM MgCl_2_, 1 mM DTT, 200 μM GMPPCP and the detergent octyl-glucopyranoside (Anatrace) at 0.2% final concentration). The filaments were observed by screening in negative stain or vitrified for cryo-EM. For vitrification, the sample was applied to Quantifoil holey carbon grids (R2/2) using a Vitrobot Mark III with 3.5 μl sample, 5 seconds blotting time and a 0 mm offset at 19°C and 100% humidity. Images were collected on an FEI T30 Polara operating at 300kV at a magnification of 31000X. Images were recorded on a Gatan K2 summit camera in super resolution mode that had a binned pixel size of 1.22 Å/pixel. The images were dose fractionated, containing 30-40 frames with a total exposure time of 6-8 seconds, 0.2 seconds per frame and a per frame dose of 1.42 electrons/Å^2^. SerialEM was used to automate data collection^7^. The defocus range was 0.8-3 μm under focus. The data was motion corrected and dose-weighted using UCSF Motioncorr2^8^. CTF parameter estimation on the non-dose-weighted but motion-corrected stacks was done using CTFFIND4 and GCTF^9,10^.

Filaments were boxed using the program e2helixboxer.py from the EMAN2 suite^11^. Particle coordinates were used to extract discrete particles using RELION 1.3-1.4^12^ and all further processing was done within the RELION suite. Multiple rounds of 2D classification identified the most well-ordered segments, and 3D autorefine was run using a Relion1.2 version with the IHRSR algorithm implemented^13,14^. The consensus helical structure was used to classify the particles without refining helical symmetry (using RELION 1.4), resulting in 2 major classes that differed slightly in rise and twist (Extended data figure 4). Particles from each class were selected and independently refined again with helical RELION 1.2 and IHRSR. Analysis of these reconstructions revealed that each structure was comprised of three linear filaments that bundle together to form a structure that resembled a triangle in cross-section (Extended data figures 3–4). The vertices of the triangle are formed through asymmetric interactions between the G-domains in adjacent filaments. The significance of these asymmetric G domain interactions has not been evaluated. The triangular arrangement of the bundled helices is unlikely to correspond to a biologically meaningful architecture, and this structure cannot form if the MID49 receptor is embedded in the outer mitochondrial membrane.

To further improve signal-to-noise, each of the three filaments in each independent half-map was segmented, extracted, resampled on a common grid and summed using UCSF Chimera^15–19^. The respective symmetrized but unfiltered half maps from each class were again aligned to a common grid, summed, averaged along the C2 symmetry axis of the DRP1 dimer. In a last step, relion_postprocess was used to add the resulting and fully symmetrized half maps (Extended data figure 4). These half maps and the final summed map, with differential B-factor sharpening per region (Extended data figures 4–9), were used for atomic modeling using Rosetta as described below.

For the projection structure of the DRP1_G362D_ rings, 2 μM protein was mixed 1:1 molar ratio with MID49_126-454_ and was allowed to sit at room temperature for an hour. The mixture was dialyzed against the assembly buffer (without detergent) overnight. The sample was collected after 12 hours and vitrified using ultra-thin 3 nm carbon support films (Ted Pella). For vitrification, a Mark III vitrobot was used with 3.5 μl sample, 0 mm offset, 100% humidity and 3.5 seconds blot time. The images were collected using an FEI TF20 microscope and SerialEM for automated data collection. The data were recorded with a Gatan K2 camera operating in super resolution mode to collect dose fractionated movie stacks using a binned pixel size of 1.234 Å/pixel. 40 frames were collected per stack (0.2 seconds per frame and 1.42 electrons/Å^2^). The movie stacks were motion corrected and the parameters of the transfer function were estimated as described above. Approximately 2000 particles were picked manually for initial 2D classification in RELION 1.4 and these averages were used as templates for further particle picking by Gautomatch (http://www.mrc-lmb.cam.ac.uk/kzhang/). Final 2D averaging of the entire rings versus quarter segments of the rings were computed using Relion1.4.

#### Model building

The general procedure for atomic model interpretation and validation using Rosetta were performed as described^20^. To obtain an initial model for DRP1, the crystal structure of nucleotide-free DRP1 (PDB ID: 4BEJ)^5^ was used for the stalk region and DRP1 G domain-BSE structures bound to GMPPCP (PDB ID: 3W6O) were used for the G-domain and BSE regions. Density-guided model completion for DRP1 was carried out with RosettaCM^21^ using this hybridization of DRP1 crystal structures. A converged solution appeared from the low-energy ensemble of the complete models generated by RosettaCM. However, among the low-energy ensemble, residues 503-612 were found to be extremely flexible without cryoEM density constraints and therefore were omitted for further coordinate refinement. For MID49, the highly homologous mouse MID49 crystal structure (81.3% identity, PDB ID: 4WOY, Extended Data Fig. 7)^6^ was used to generate a homology model using RosettaCM and used as the starting model.

To enable fragment-based, density-guided model refinement with missing residues (503-612, DRP1), Rosetta iterative local rebuilding tool was customized to disallow backbone rebuilding at breaks within a single chain. Multiple rounds of refinement were done for each component against one half map (training map), and the other half map (validation map) was used to monitor overfitting according to the detailed procedure described in Wang et al.^20^.

With the refined model of DRP1 and MID49, we further refined the model in the context of a full assembly that included 8 identical copies of each protein, Mg^2+^ and nucleotide which included all possible inter-domain molecular interactions in the filament (Extended data figure 8).

Pseudo-symmetry was used^22^ to enable and facilitate the energy evaluations of all neighboring interactions around the asymmetric unit (Green model, shown in Extended data figure 8) for final model refinement of the full assembly. To this end, refinement was done against the training map. Finally, the half maps were used to determine a weight for the density map that did not introduce overfitting. Using the weight and with the symmetry imposed, the whole assembly of DRP1 and MID49 was refined in the full map, followed by B-factors refinement^23^. Finally, quantification of buried surface area and the number and nature of the bonds involved for each DRP1-MID49 interaction interface modeled by Rosetta were performed with the PISA server (http://www.ebi.ac.uk/pdbe/pisa/).

To design a molecular model for the closed 12-dimer DRP1 rings, we used the diameter, thickness, and angles revealed by the 2D cryoEM class averages of the DRP1_G362D_ rings stabilized with GMPPCP. The atomic coordinates determined above using RosettaCM were used to build the ring in sections, first with repeating dimers of the interface-2 “X-shaped” stalk, then the BSE and finally the G-domains and the angles between these sections were iteratively adjusted until calculated projections of the molecular model corresponded with the features of the experimental projection densities. Both the top (Fig 4b-c) and the side view (Extended data Figure 15) were used as constraints. The complete atomic model of ring was finally refined in Phenix^24^ to minimize clashes.

**Extended data Table 1.**
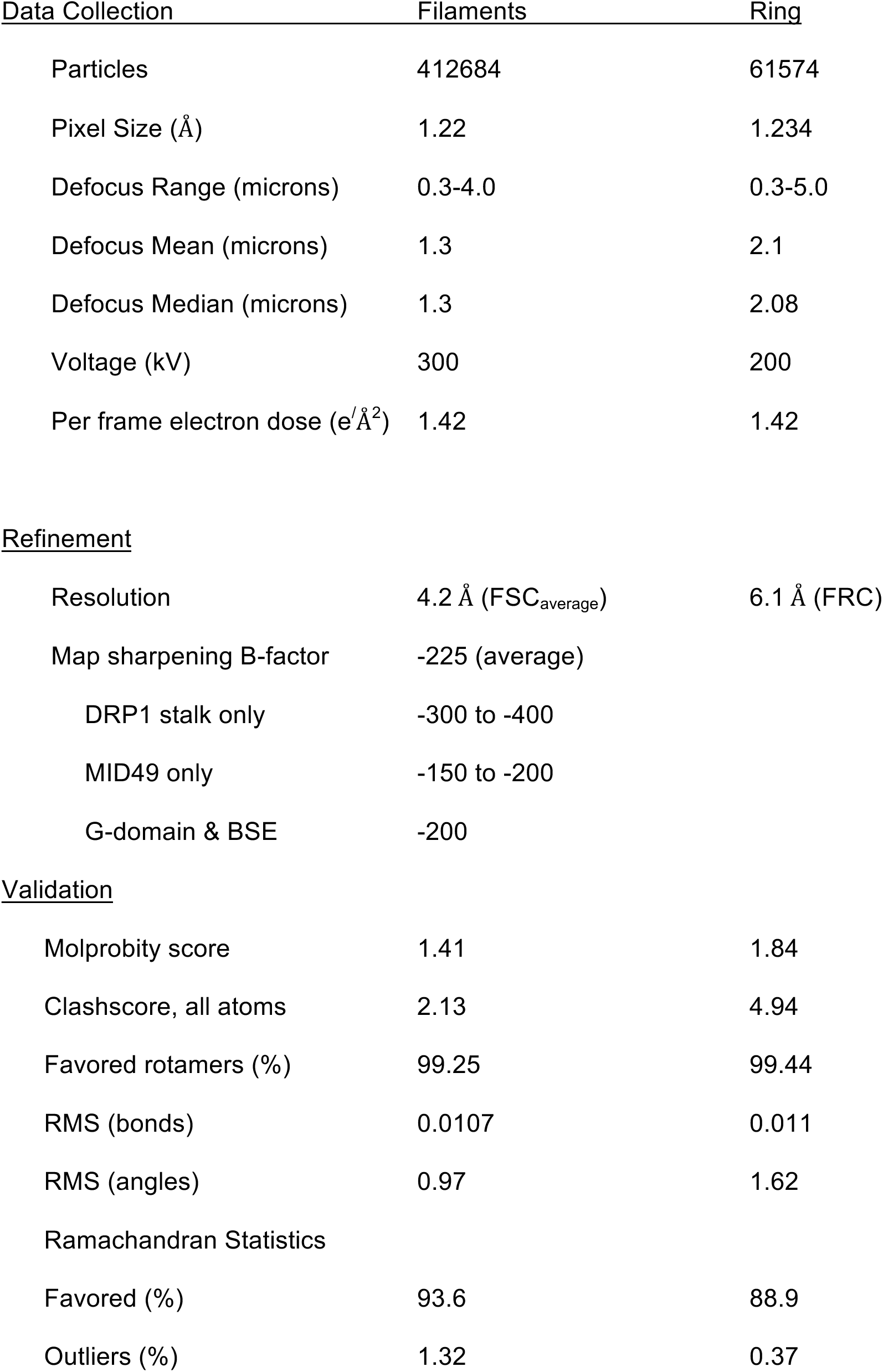

## Extended data figures

**Extended data figure 1:**
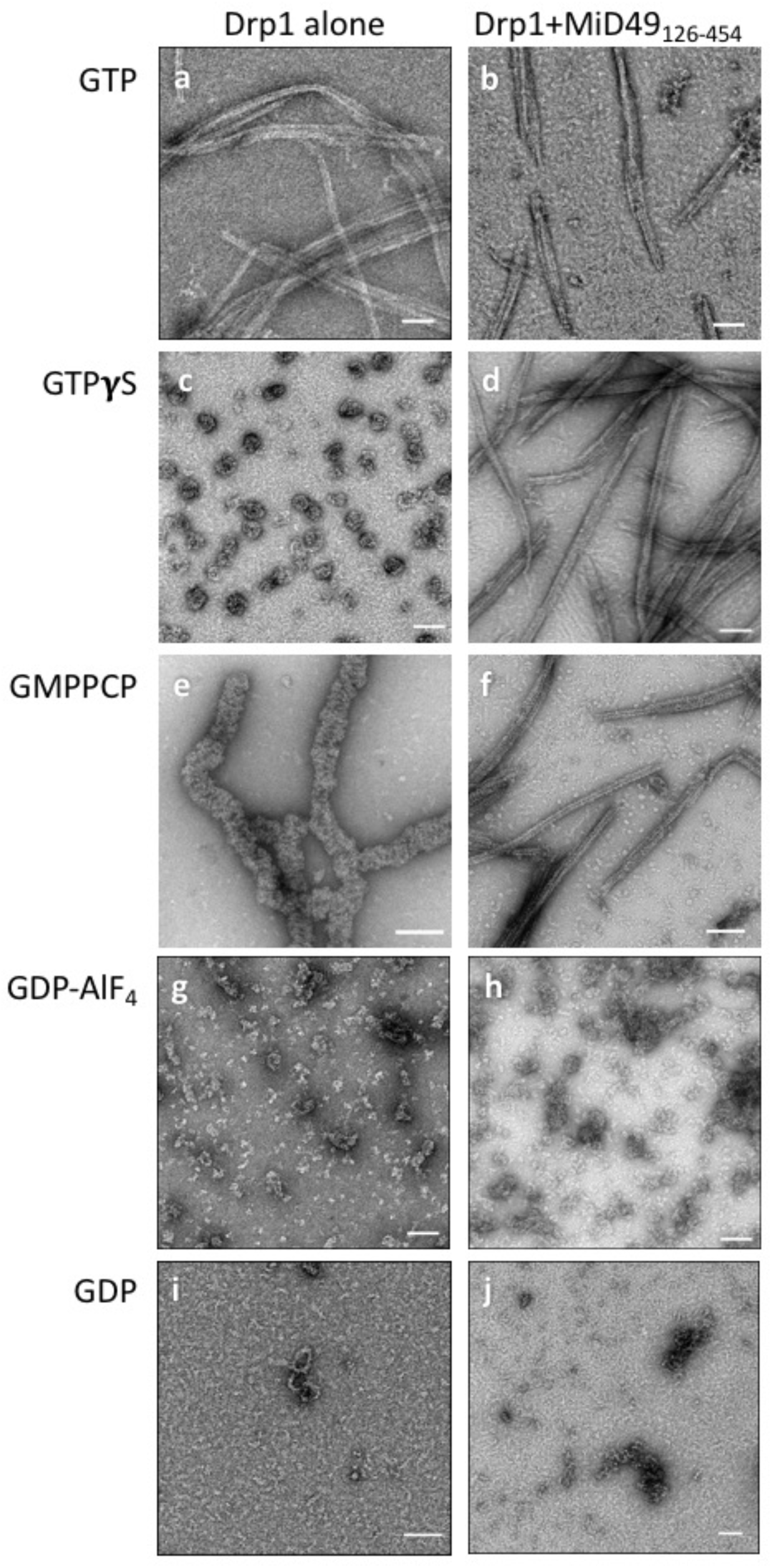
DRP1 assembly states visualized with negative stain electron microscopy in the presence of different guanine nucleotides and stoichiometric MID49_126-454_ ([2μM] for both proteins). Bars = 100nm.

**Extended data figure 2:**
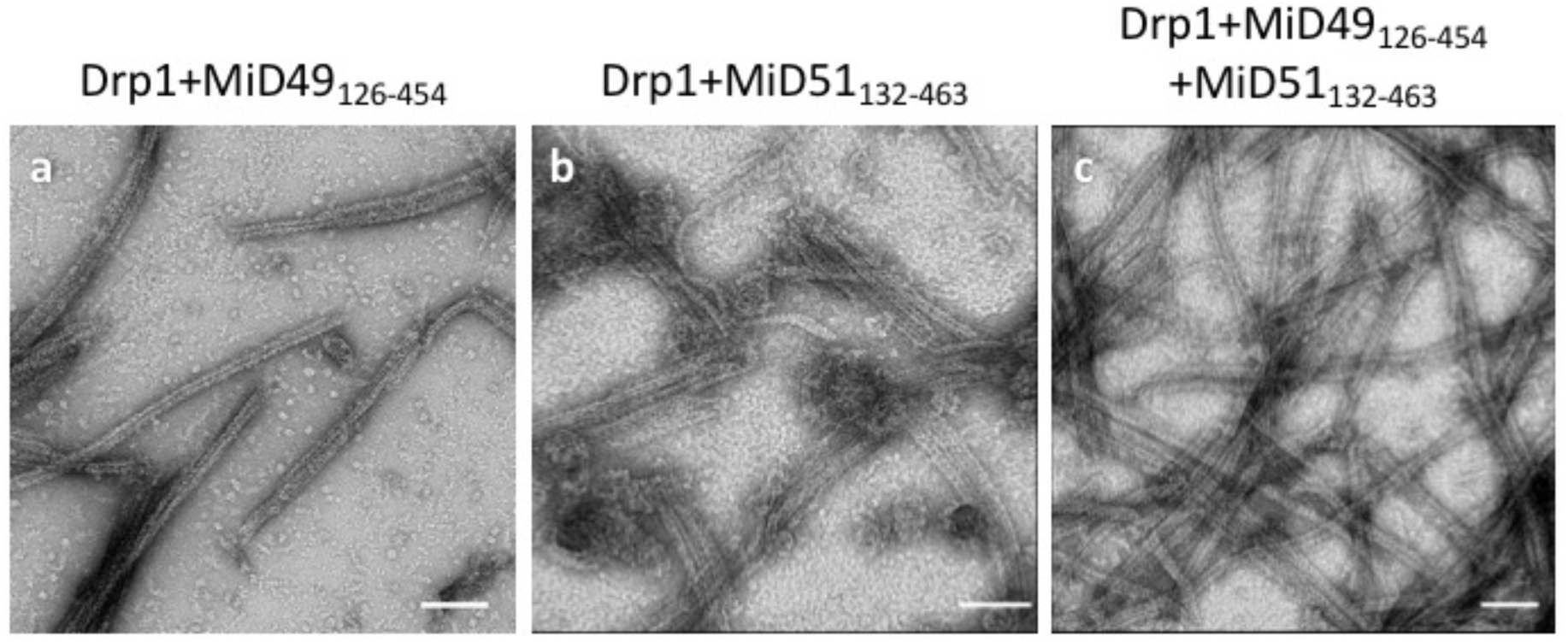
MID49_126-454_ and MID51_132-463_ form indistinguishable assemblies with DRP1 by negative stain electron microscopy. **(a)** DRP1 plus MID49_126-454_ and GMPPCP; **(b)** DRP1 plus MID51_132-463_ and GMPPCP, **(c)** DRP1 plus both MID49 and MID51. Bars = 100nm.

**Extended data figure 3:**
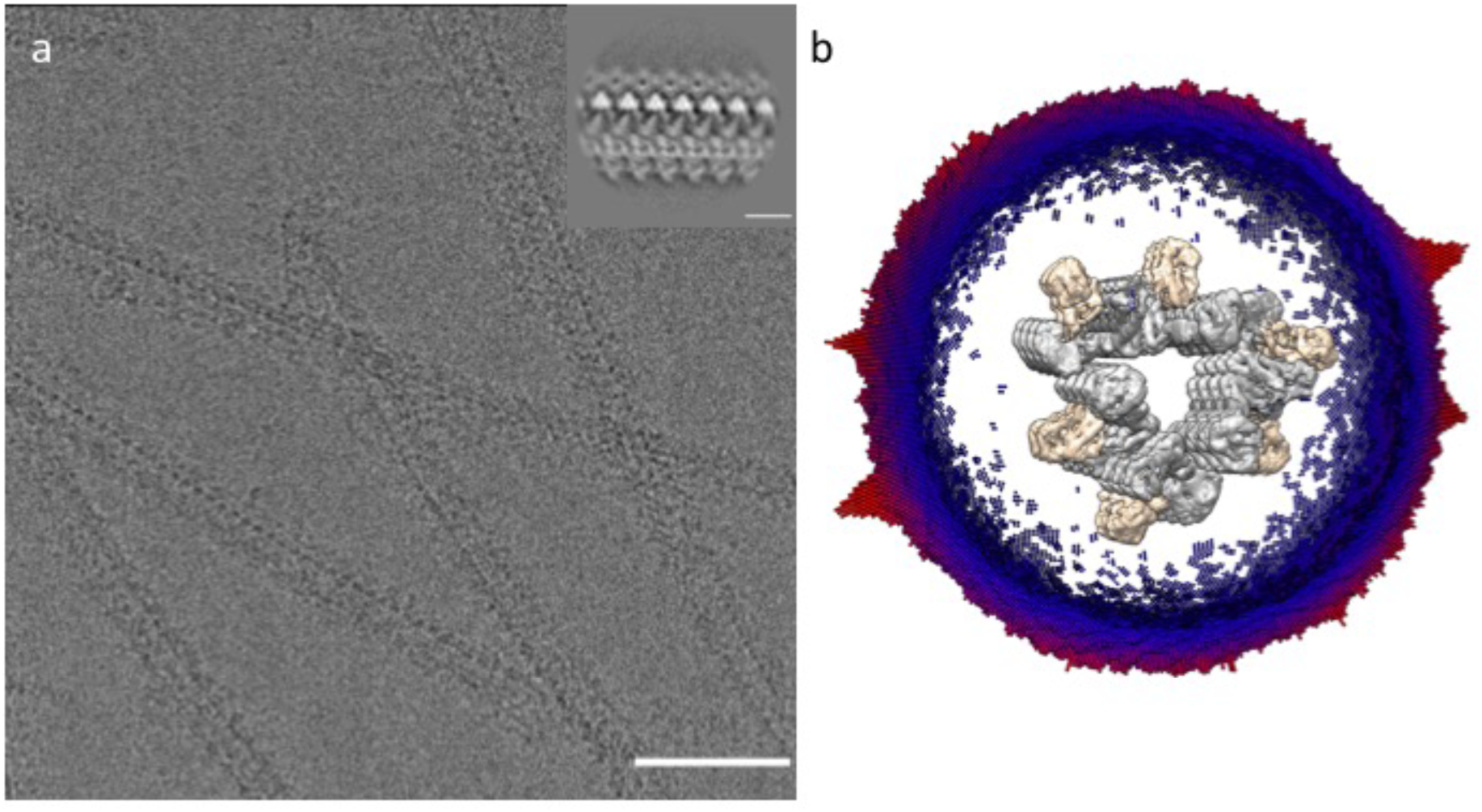
CryoEM imaging and reconstruction. **(a)** An electron cryo-micrograph of DRP1-MID49_126-454_ filaments formed with GMPPCP. Inset shows a representative 2D class average. Bar =100nm, Inset bar =10nm. **(b)** Oblique cross-section of the 3D reconstruction and the distribution of views determined during helical reconstruction. DRP1 density is rendered in grey, MID49 in golden yellow.

**Extended data figure 4:**
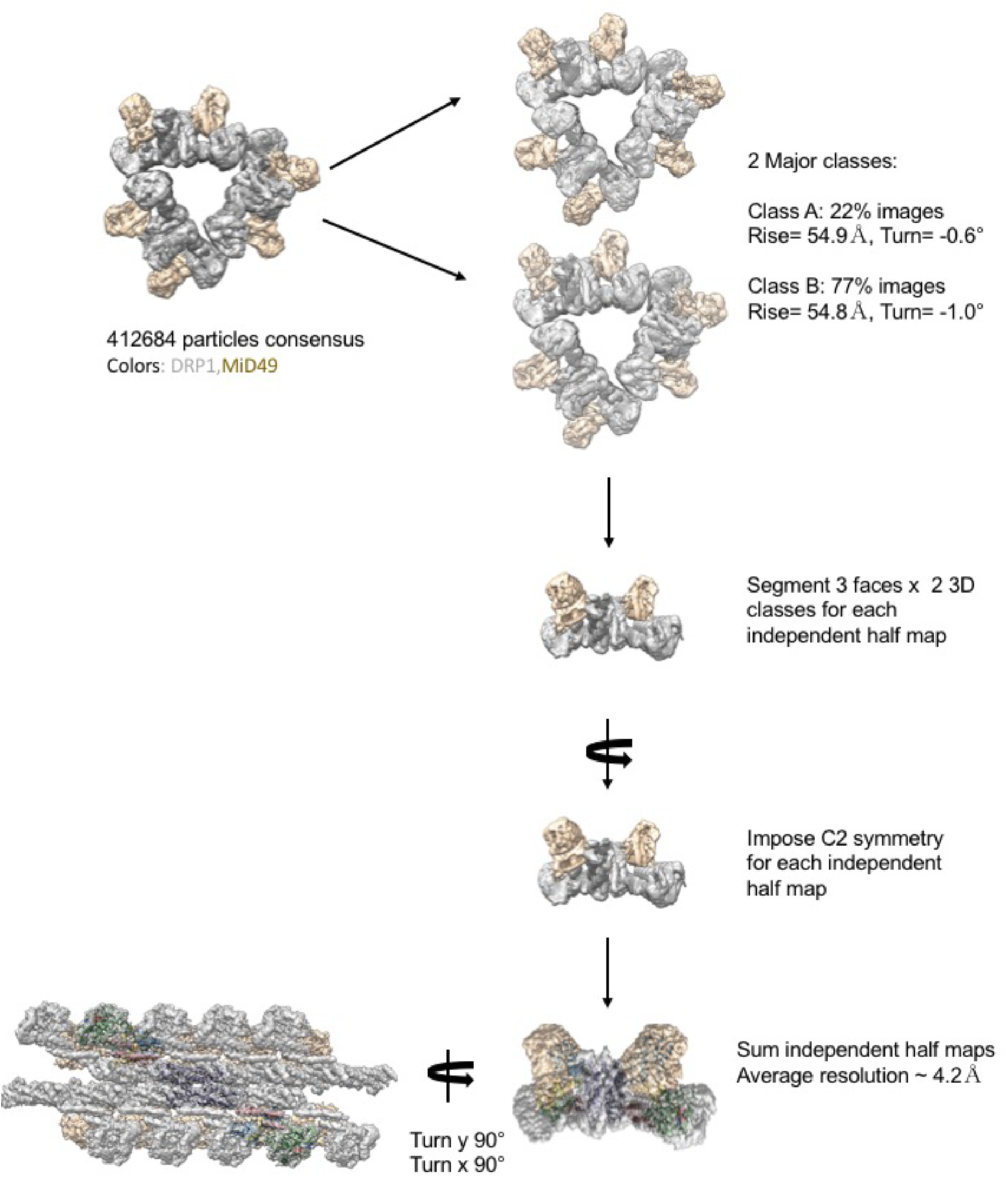
Raw particle numbers and workflow for the reconstruction protocol and imposition of symmetries. DRP1 density is rendered in grey, MID49 in golden yellow.

**Extended data figure 5:**
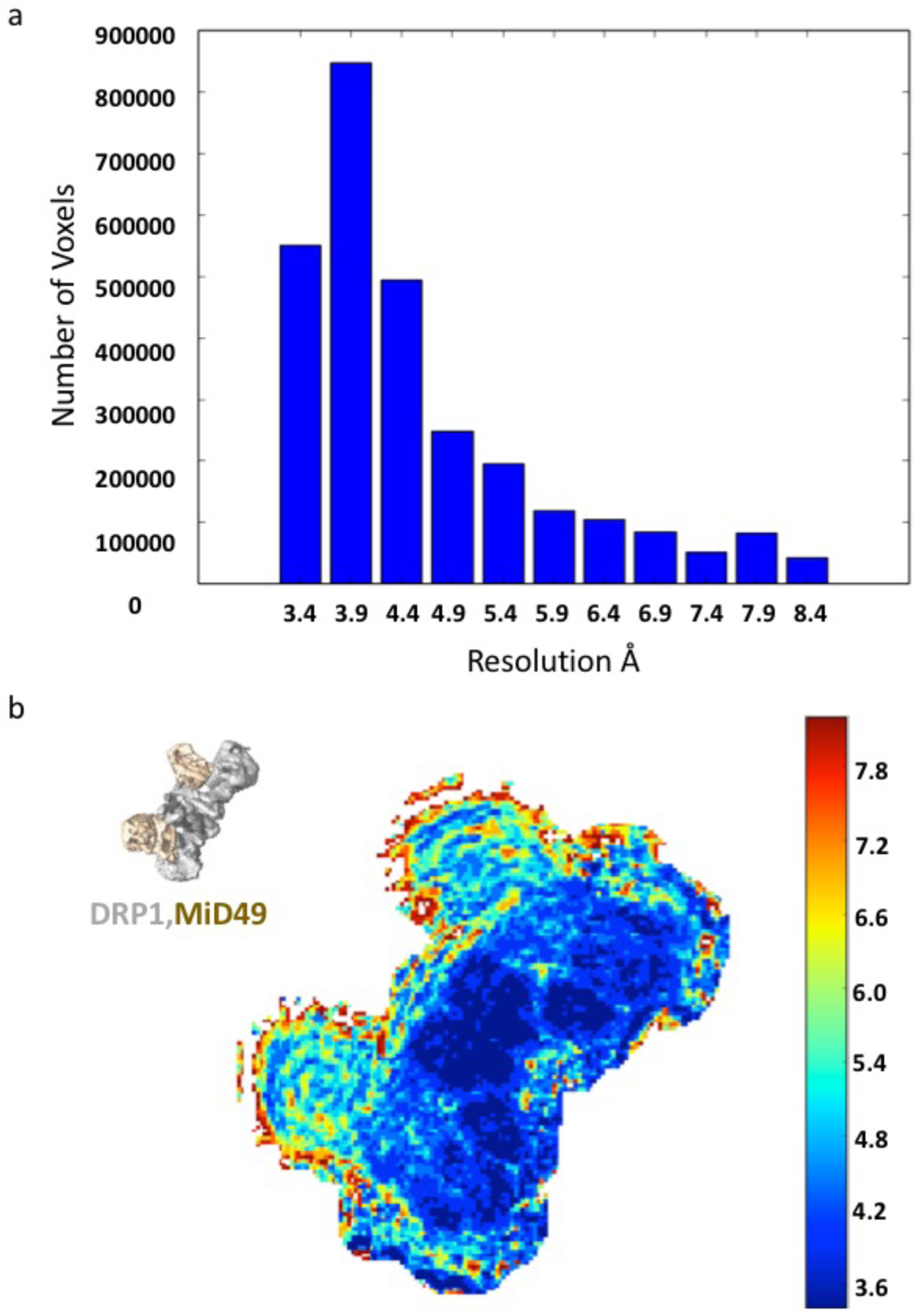
Local resolution estimates computed by Resmap^25^. **(a)** Histogram of voxels values, and **(b)** Heat map of local resolution estimates displayed for a cross-section through the reconstruction.

**Extended data figure 6:**
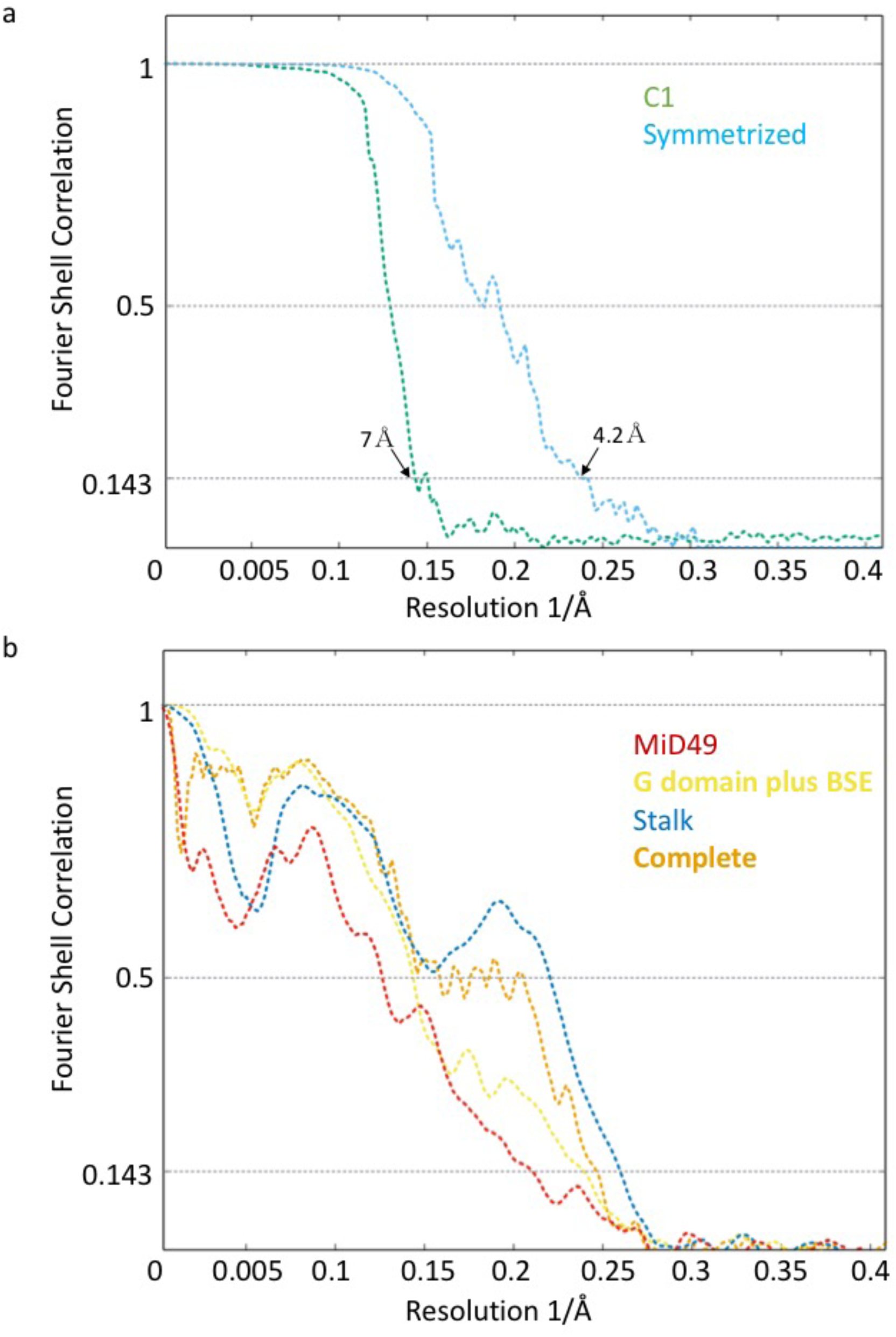
Fourier Shell Correlation plots for **(a)** the half-maps with and without imposed symmetry; and **(b)** model-to-map correlations for the averaged as well as each subregion of the structure using the atomic coordinates and B-factors determined using Rosetta.

**Extended data figure 7:**
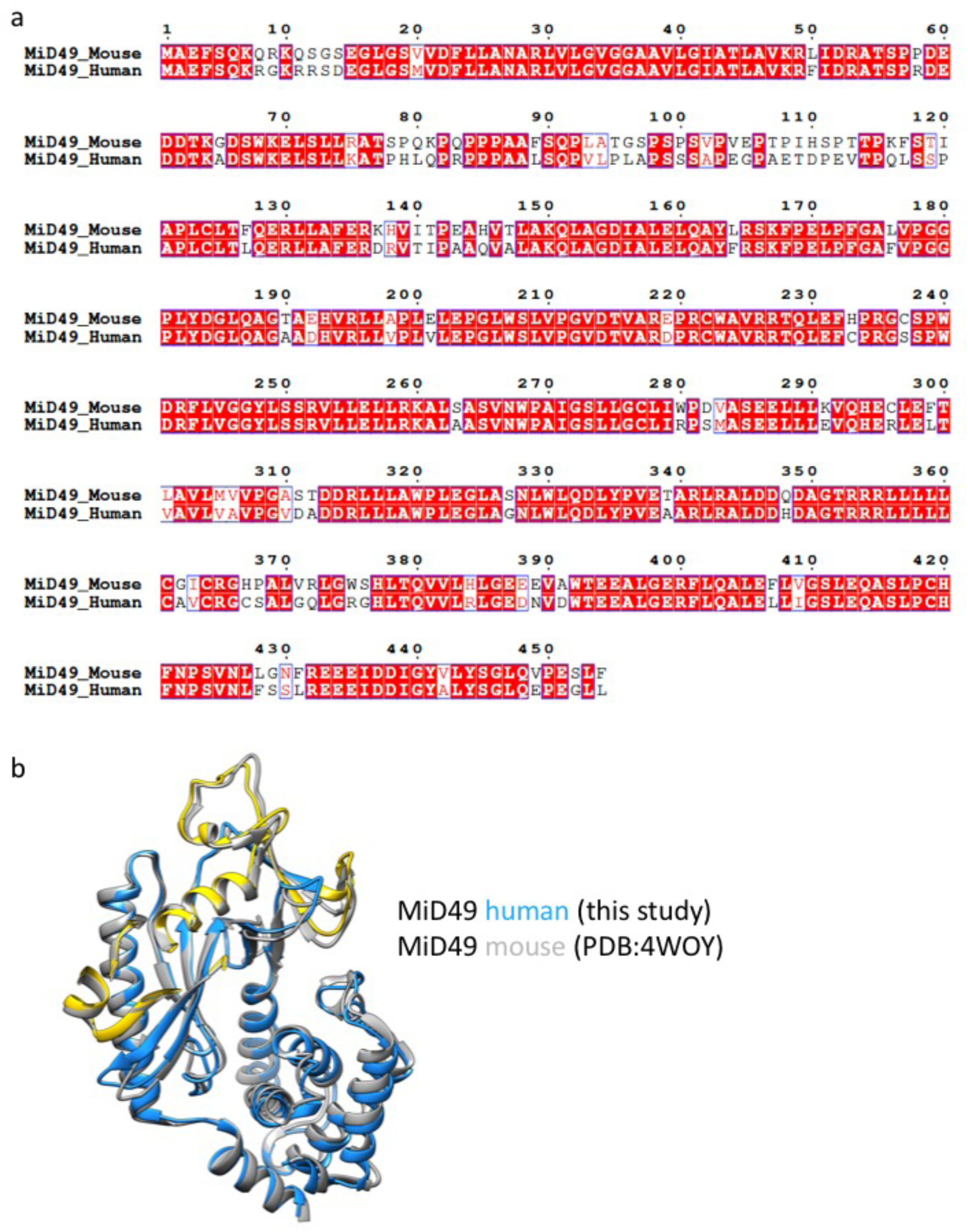
Modeling of human MID49. **(a)** Sequence alignment between human and mouse MID49 sequences. **(b)** Overlay of the homology model of human MID49_126-454_ (blue, with yellow DRR, ribbon) modeled within the cryoEM density versus the mouse MID49 crystal structure (PDB ID: 4WOY, grey ribbon)^6^.

**Extended data figure 8:**
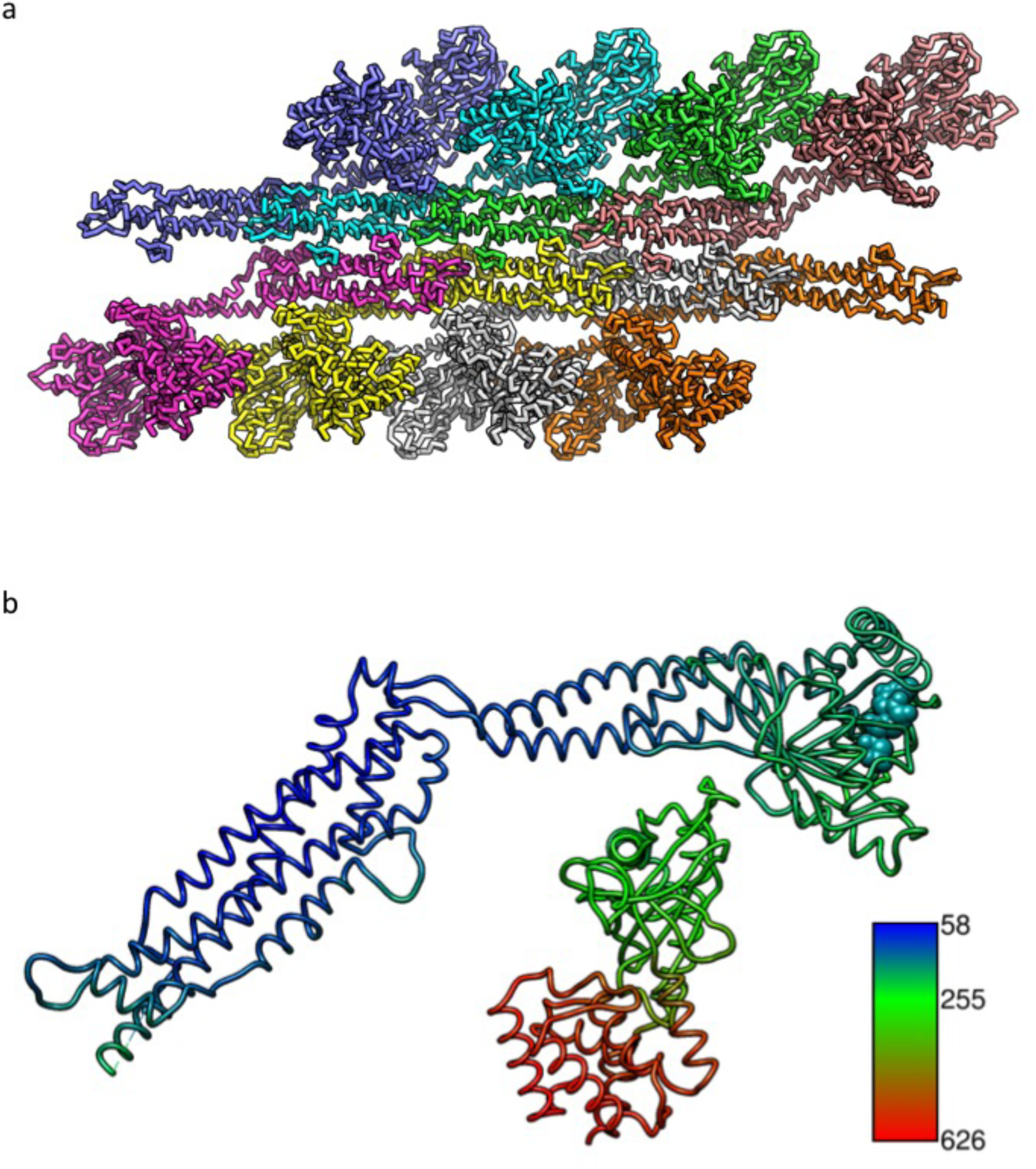
Rosetta refinement. **(a)** Symmetric unit of the filament with 8 chains of DRP1 and 8 chains of MID49 refined using Rosetta to enforce symmetry and account for all possible inter-molecular interfaces. **(b)** Atomic B-factors for the DRP1 and MID49 models (ribbon) and bound GMPPCP (space filling).

**Extended data figure 9:**
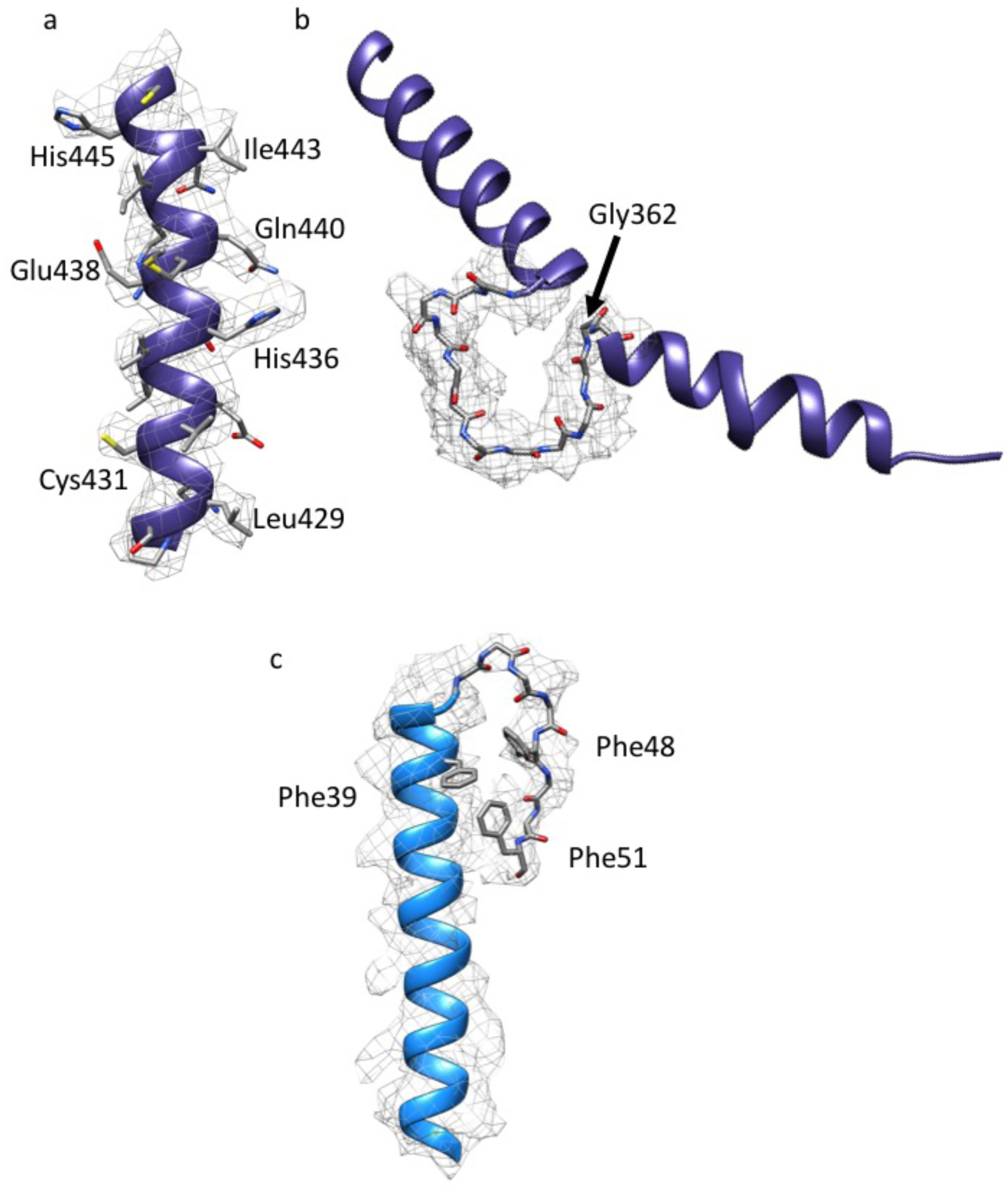
Examples of model fitting to B-factor sharpened density for **(a)** a helix from the DRP1 stalk, **(b)** the backbone of the L1N^S^ loop, and **(c)** a helix and loop from MID49.

**Extended data figure 10:**
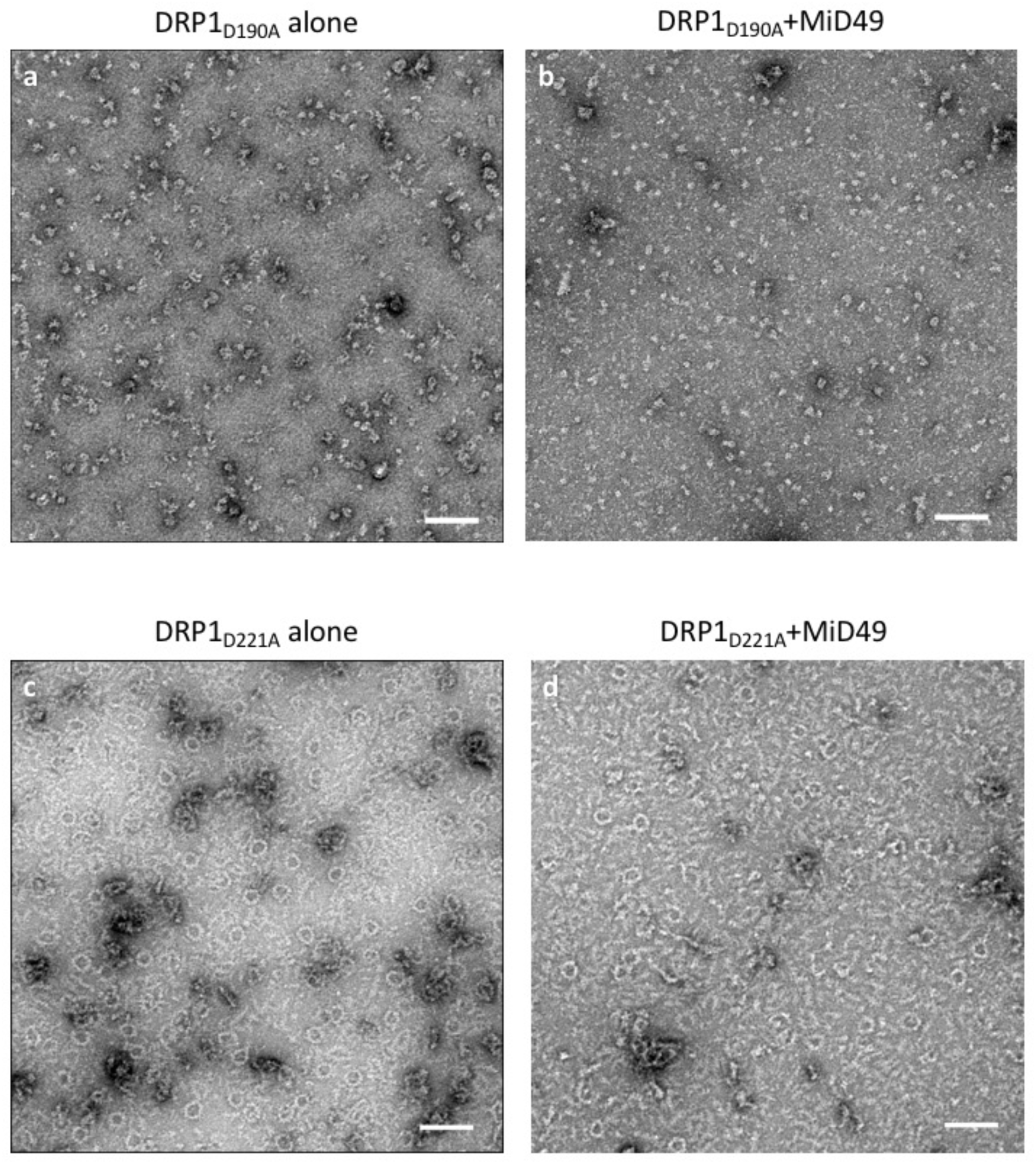
DRP1 assembly and coassembly reactions with GMPPCP for **(a)** DRP1_D190A_ alone, **(b)** DRP1_D190A_+MID49, **(c)** DRP1_D221A_ alone, and **(d)** DRP1_D221A_+MID49. Bars = 100nm.

**Extended data figure 11:**
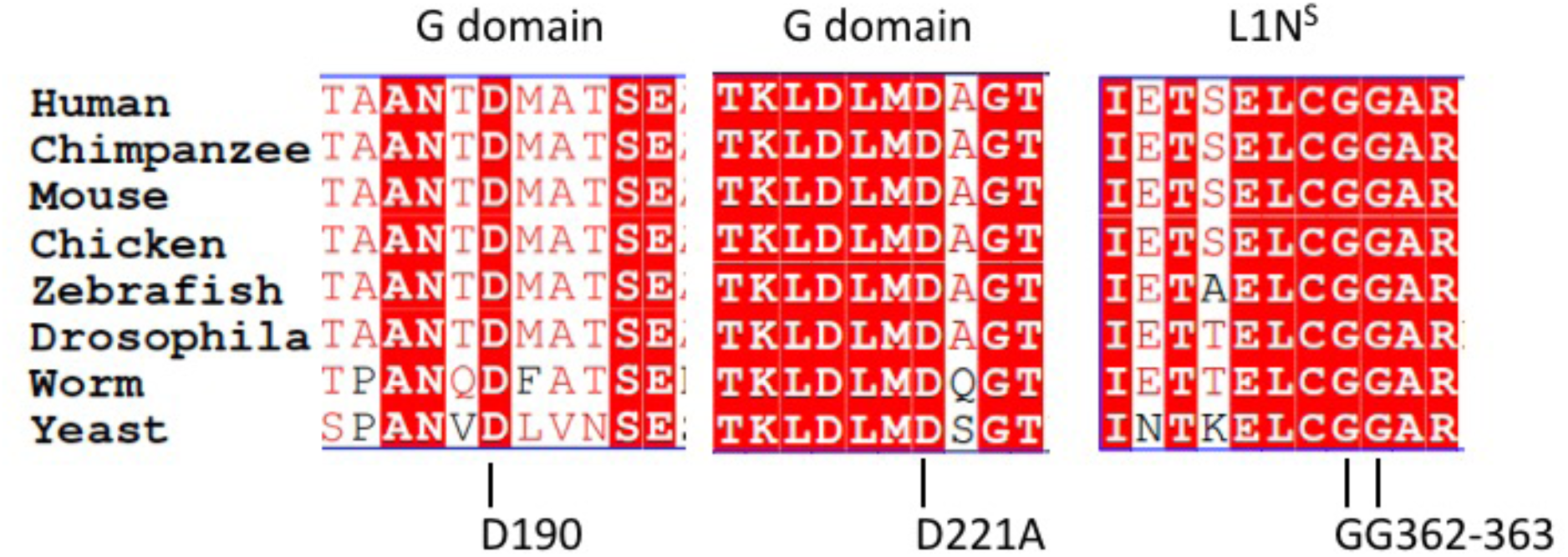
Multiple sequence alignment of the regions near and including the DRP1 residues mutated in this study: D190, D221 and G362. The residue numbers apply to human DRP1, isoform 2 (UNIPROT identifier: O00429-3 also known as DLP1a).

**Extended data figure 12:**
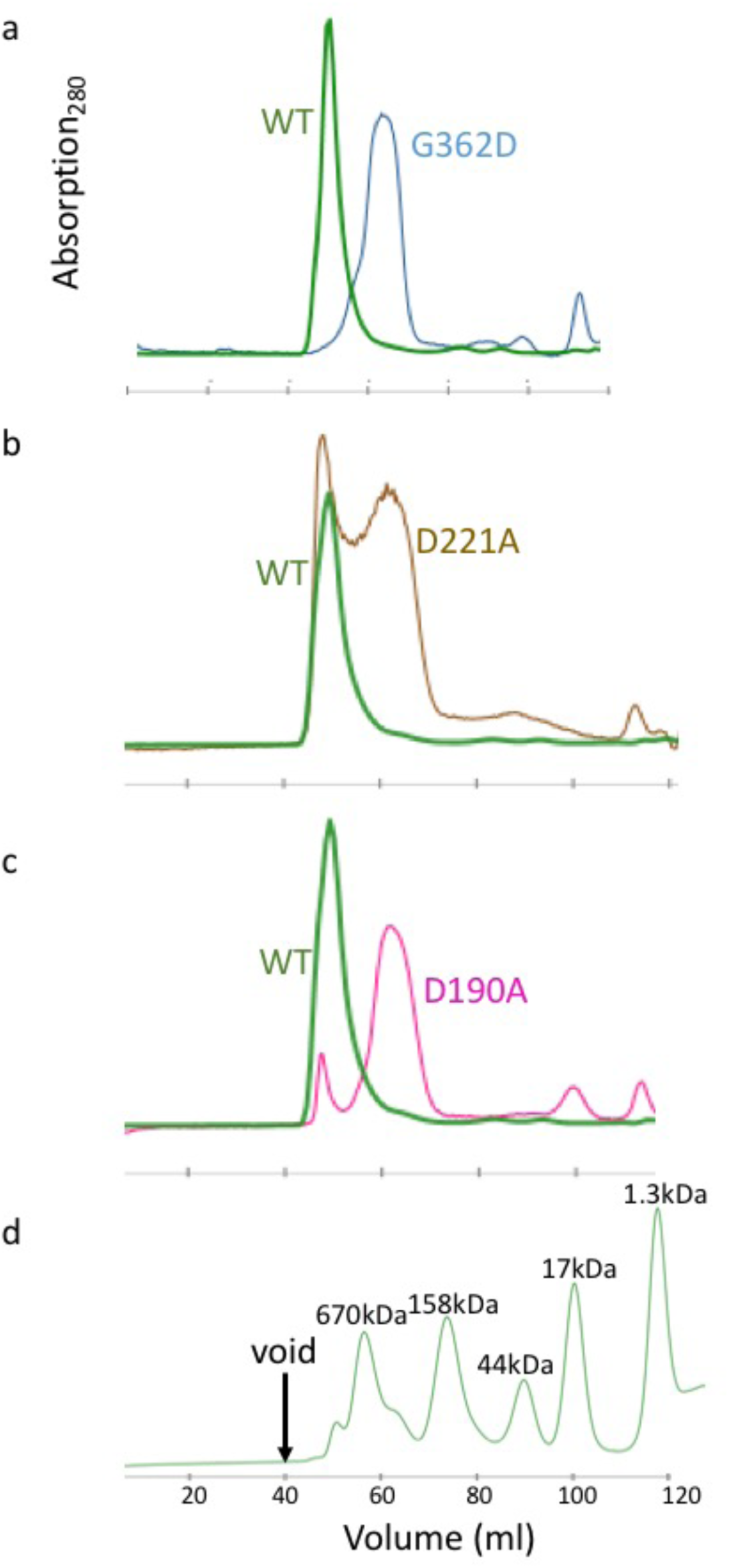
Size exclusion chromatography traces for DRP1 wild-type and mutants used in the study. **(a)** Comparison between wild type (WT) versus DRP1_G362D_, **(b)** WT versus DRP1_D221A_, and **(c)** WT versus DRP1_D190A_. (**d**) Standards for molecular weight comparison.

**Extended data figure 13:**
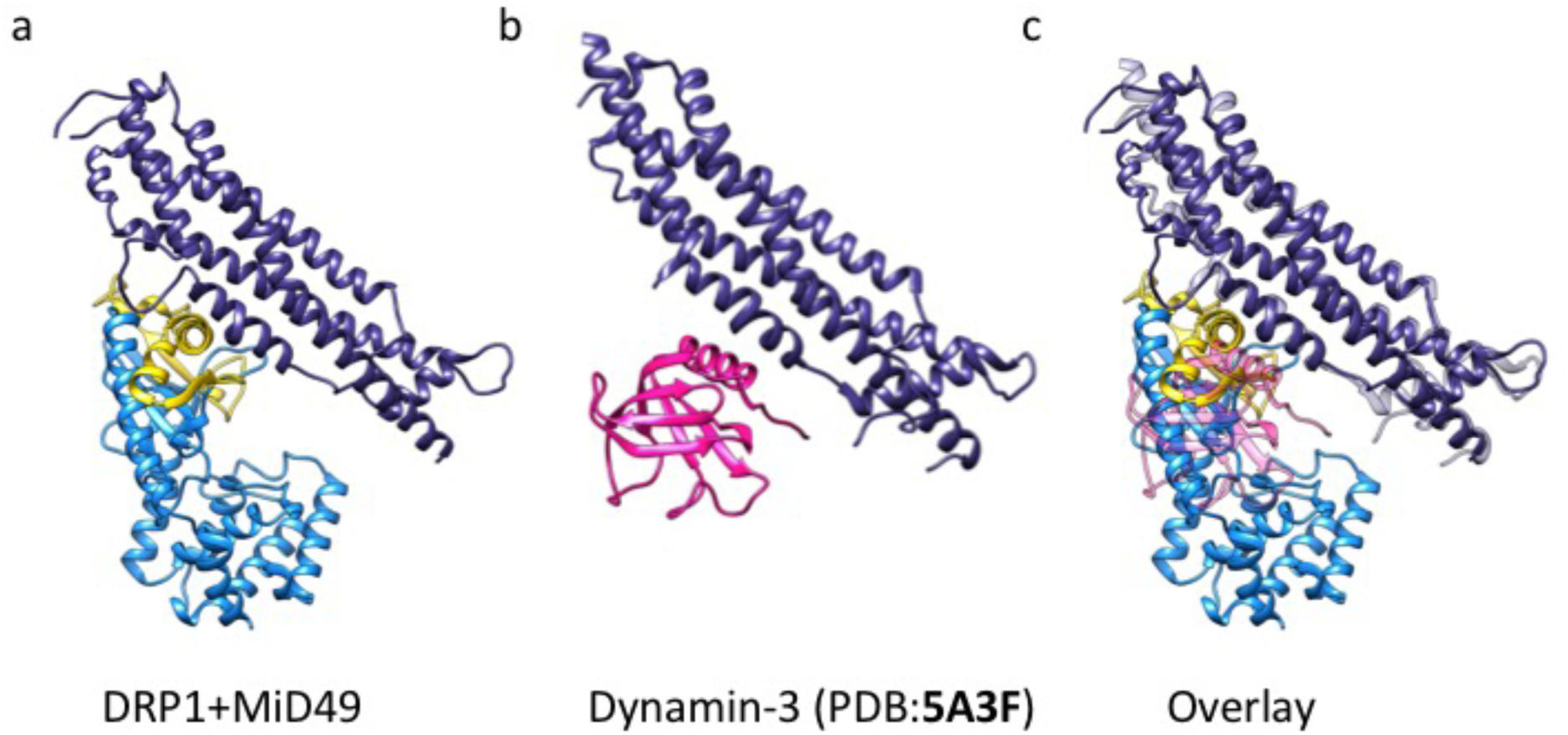
Structural similarities between the third DRP1-MID49 interaction interface that includes the L1N^S^ loop and the interaction between the Pleckstrin Homology (PH) domain and the stalk of Dynamin-3 (PDB ID:5A3F)^26^. **(a)** DRP1-MID49 interaction at MID-interaction interface 3, **(b)** the PH domain bound to the stalk of Dynamin-3, and **(c)** Overlay of a and b.

**Extended data figure 14:**
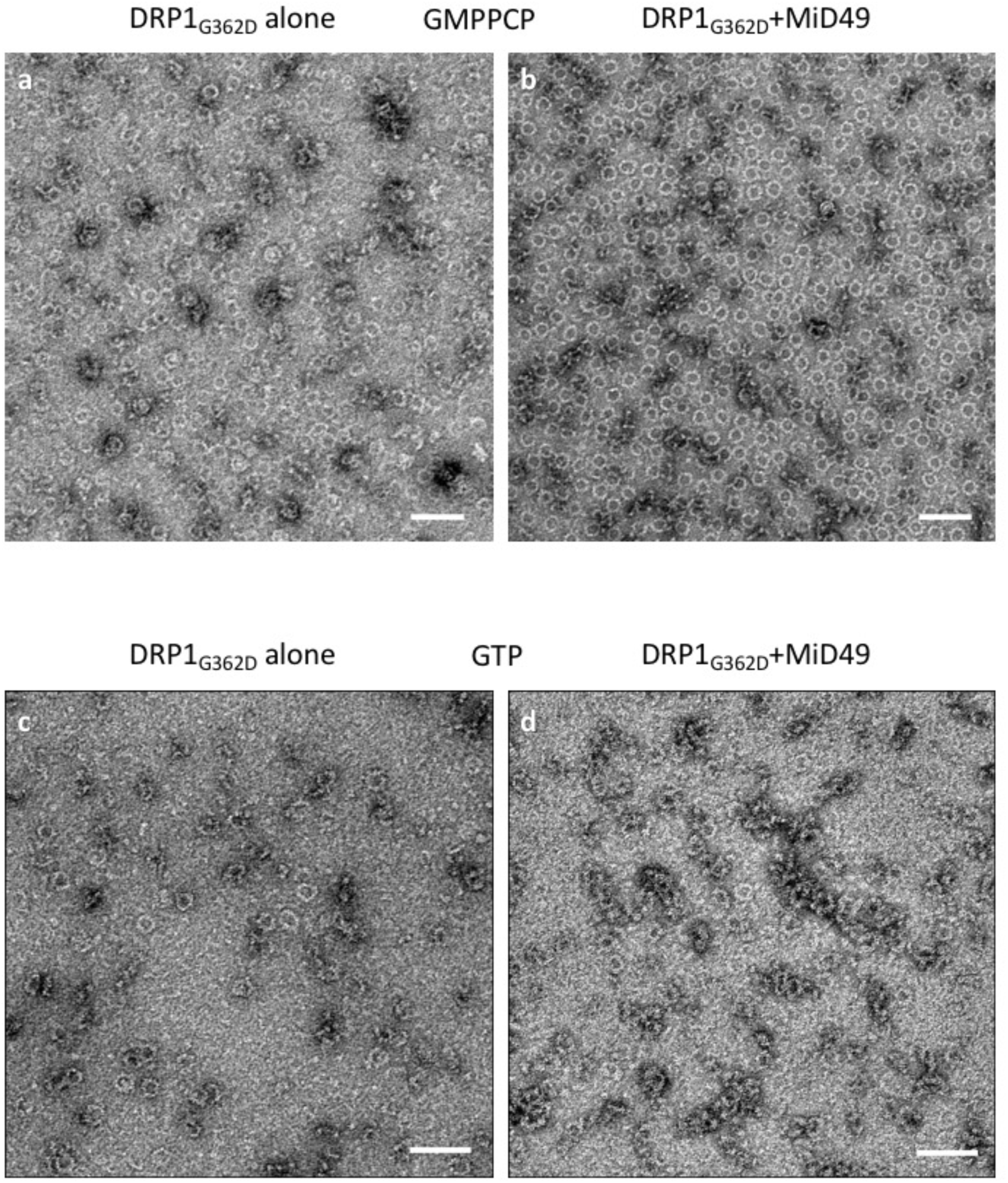
DRP1_G362D_ assembly and coassembly reactions with GMPPCP or GTP. DRP1_G362D_ forms rings but not linear filaments without MID49 **(a,c)**; and with MID49 present **(b,d)**; with GMPPCP **(a-b)**; or with GTP **(c-d)**. Bars = 100 nm.

**Extended data figure 15:**
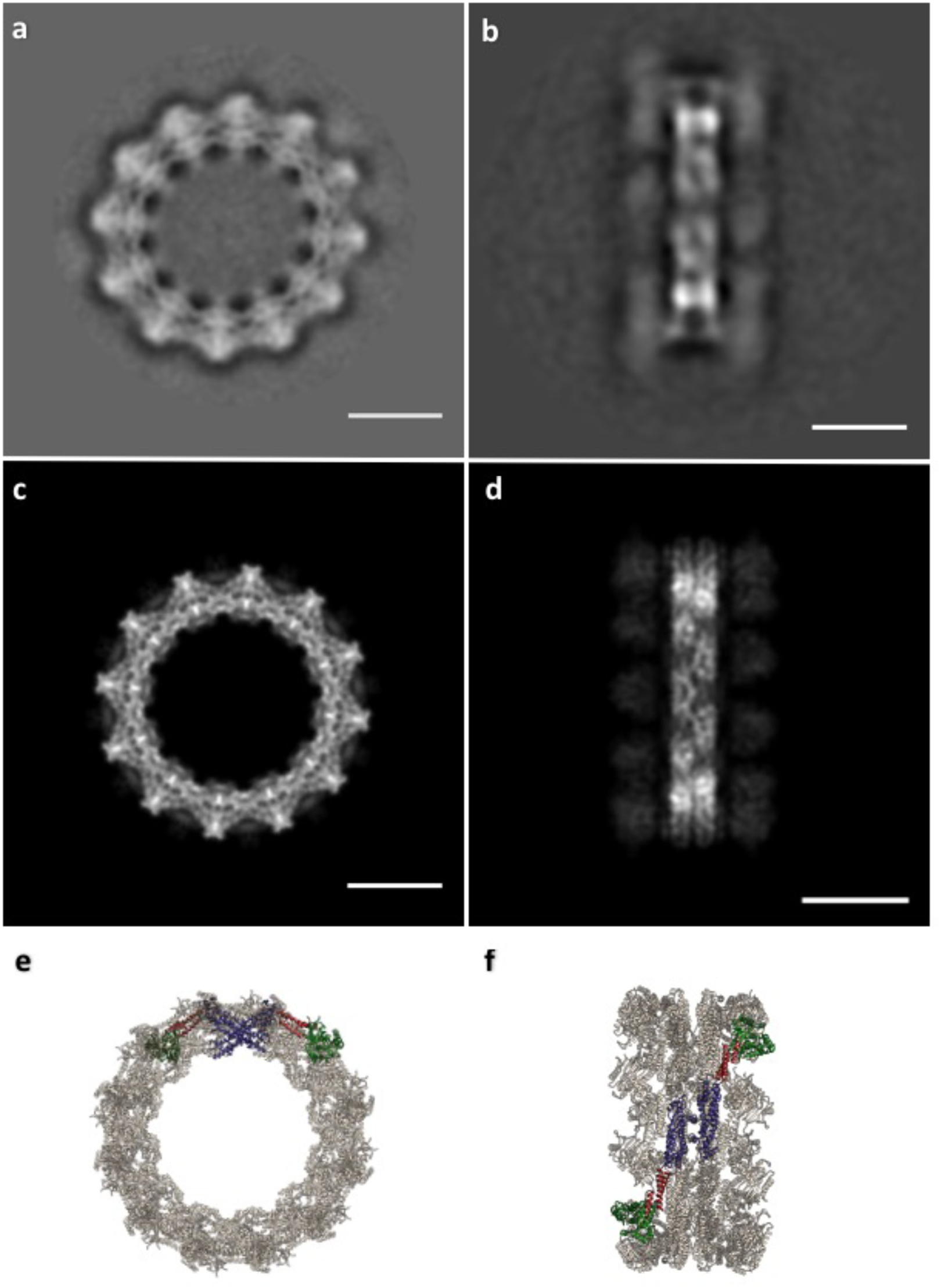
DRP1_G362D_ forms 12-dimer closed rings. **(a)** 2D class average of the rings; **(b)** 2D class average of infrequent, orthogonal or “side” views used as a constraint during model building; **(c)** “top” and **(d)** “side” projections of the model; **(e)** “top” and **(f)** “side” views of the final model rendered as ribbons. Bars =100Å. Green: G-domain, Red: <u>B</u>undle <u>S</u>ignaling <u>E</u>lement (BSE), Purple: Stalk region.

## Extended data movie Legends

**Extended data movie 1:** Nucleotide-induced conformational changes in the G-domain and Bundle Signaling Element (BSE).

**Extended data movie 2:** Nucleotide-induced conformational changes in the G-domain and Bundle Signaling Element (BSE) in the context of a full-length DRP1 dimer (“side” view).

**Extended data movie 3:** Nucleotide-induced conformational changes in the G-domain and Bundle Signaling Element (BSE) in the context of a full-length DRP1 dimer (“top” view).

**Extended data movie 4:** MID receptors engage a specific nucleotide-bound conformation of DRP1 tetramers and promote formation of a linear copolymer. Subsequent nucleotide hydrolysis and exchange promotes receptor dissociation and tetramer bending in the closed rings.

## 1 Overall quality at a glance ⓘ

The following experimental techniques were used to determine the structure:

### ELECTRON MICROSCOPY

The reported resolution of this entry is unknown.

Percentile scores (ranging between 0-100) for global validation metrics of the entry are shown in the following graphic. The table shows the number of entries on which the scores are based.

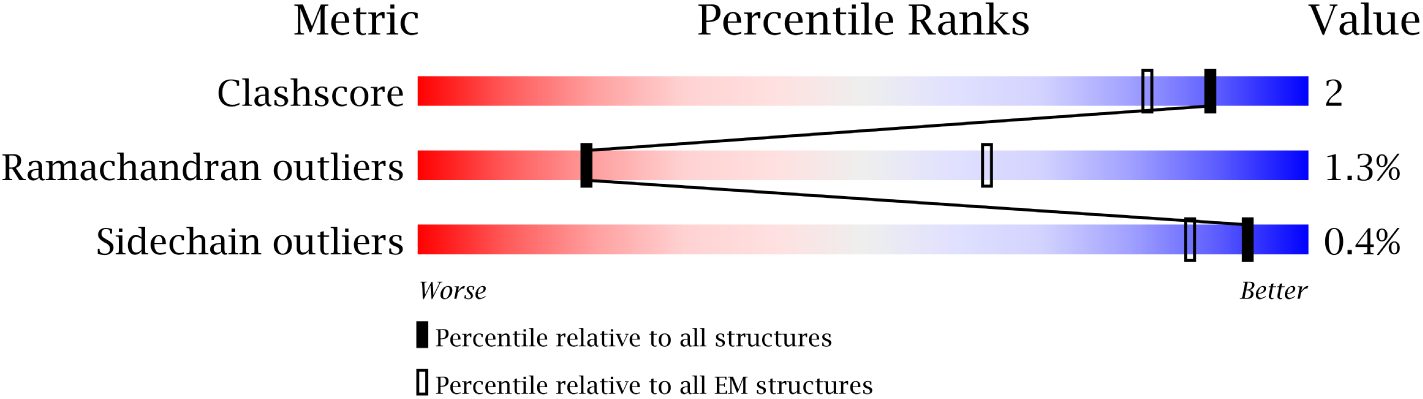

| Metric | Whole archive<br>(#Entries) | EM structures<br>(#Entries) |
| --- | --- | --- |
| Clashscore | 125131 | 1336 |
| Ramachandran outliers | 121729 | 1120 |
| Sidechain outliers | 121581 | 1026 |

The table below summarises the geometric issues observed across the polymeric chains. The red, orange, yellow and green segments on the bar indicate the fraction of residues that contain outliers for >=3, 2, 1 and 0 types of geometric quality criteria. A grey segment represents the fraction of residues that are not modelled. The numeric value for each fraction is indicated below the corresponding segment, with a dot representing fractions <=5%

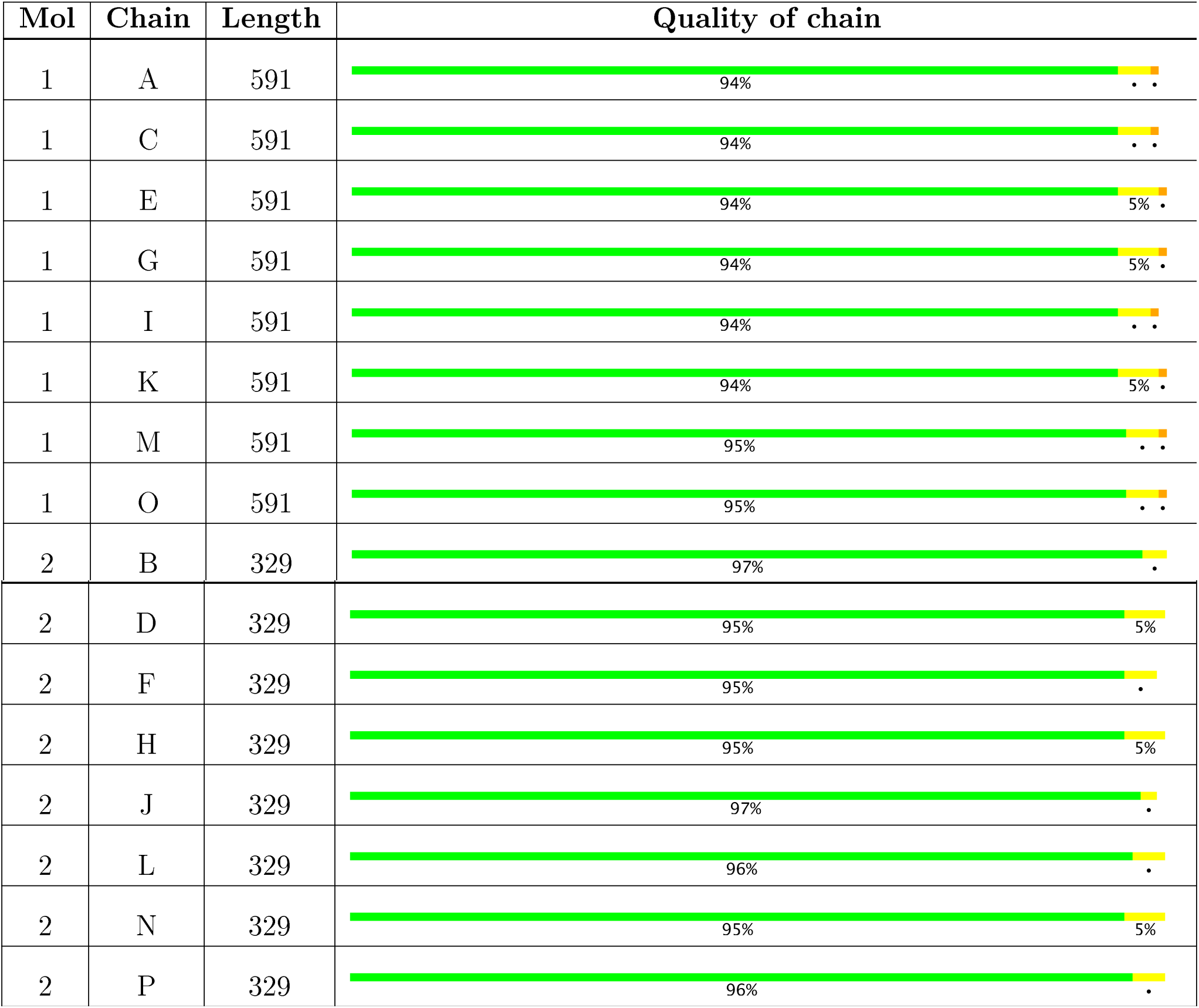

## 2 Entry composition ⓘ

There are 2 unique types of molecules in this entry. The entry contains 116960 atoms, of which 59040 are hydrogens and 0 are deuteriums.

In the tables below, the AltConf column contains the number of residues with at least one atom in alternate conformation and the Trace column contains the number of residues modelled with at most 2 atoms.

| Mol | Chain | Residues | Atoms |  |  |  |  |  |  |  | AltConf | Trace |
| --- | --- | --- | --- | --- | --- | --- | --- | --- | --- | --- | --- | --- |
| 1 | A | 591 | Total | C | H | Mg | N | O | P | S | 0 | 1 |
|  |  |  | 9493 | 2954 | 4801 | 1 | 821 | 893 | 3 | 20 |  |  |
| 1 | C | 591 | Total | C | H | Mg | N | O | P | S | 0 | 1 |
|  |  |  | 9493 | 2954 | 4801 | 1 | 821 | 893 | 3 | 20 |  |  |
| 1 | E | 591 | Total | C | H | Mg | N | O | P | S | 0 | 1 |
|  |  |  | 9493 | 2954 | 4801 | 1 | 821 | 893 | 3 | 20 |  |  |
| 1 | G | 591 | Total | C | H | Mg | N | O | P | S | 0 | 1 |
|  |  |  | 9493 | 2954 | 4801 | 1 | 821 | 893 | 3 | 20 |  |  |
| 1 | I | 591 | Total | C | H | Mg | N | O | P | S | 0 | 1 |
|  |  |  | 9493 | 2954 | 4801 | 1 | 821 | 893 | 3 | 20 |  |  |
| 1 | K | 591 | Total | C | H | Mg | N | O | P | S | 0 | 1 |
|  |  |  | 9493 | 2954 | 4801 | 1 | 821 | 893 | 3 | 20 |  |  |
| 1 | M | 591 | Total | C | H | Mg | N | O | P | S | 0 | 1 |
|  |  |  | 9493 | 2954 | 4801 | 1 | 821 | 893 | 3 | 20 |  |  |
| 1 | O | 591 | Total | C | H | Mg | N | O | P | S | 0 | 1 |
|  |  |  | 9493 | 2954 | 4801 | 1 | 821 | 893 | 3 | 20 |  |  |

- Molecule 2 is a protein.
| Mol | Chain | Residues | Atoms |  |  |  |  |  | AltConf | Trace |
| --- | --- | --- | --- | --- | --- | --- | --- | --- | --- | --- |
| 2 | B | 329 | Total<br>5127 | C<br>1630 | H<br>2579 | N<br>445 | O<br>465 | S<br>8 | 0 | 0 |
| 2 | D | 329 | Total<br>5127 | C<br>1630 | H<br>2579 | N<br>445 | O<br>465 | S<br>8 | 0 | 0 |
| 2 | F | 329 | Total<br>5127 | C<br>1630 | H<br>2579 | N<br>445 | O<br>465 | S<br>8 | 0 | 0 |
| 2 | H | 329 | Total<br>5127 | C<br>1630 | H<br>2579 | N<br>445 | O<br>465 | S<br>8 | 0 | 0 |
| 2 | J | 329 | Total<br>5127 | C<br>1630 | H<br>2579 | N<br>445 | O<br>465 | S<br>8 | 0 | 0 |
| 2 | L | 329 | Total<br>5127 | C<br>1630 | H<br>2579 | N<br>445 | O<br>465 | S<br>8 | 0 | 0 |
| 2 | N | 329 | Total<br>5127 | C<br>1630 | H<br>2579 | N<br>445 | O<br>465 | S<br>8 | 0 | 0 |

| Mol | Chain | Residues | Atoms |  |  |  |  |  | AltConf | Trace |
| --- | --- | --- | --- | --- | --- | --- | --- | --- | --- | --- |
|  |  |  | Total | C | H | N | O | S |  |  |
| 2 | P | 329 | 5127 | 1630 | 2579 | 445 | 465 | 8 | 0 | 0 |

## 3 Residue-property plots ⓘ

These plots are drawn for all protein, RNA and DNA chains in the entry. The first graphic for a chain summarises the proportions of the various outlier classes displayed in the second graphic. The second graphic shows the sequence view annotated by issues in geometry. Residues are colorcoded according to the number of geometric quality criteria for which they contain at least one outlier: green = 0, yellow = 1, orange = 2 and red = 3 or more. Stretches of 2 or more consecutive residues without any outlier are shown as a green connector. Residues present in the sample, but not in the model, are shown in grey.

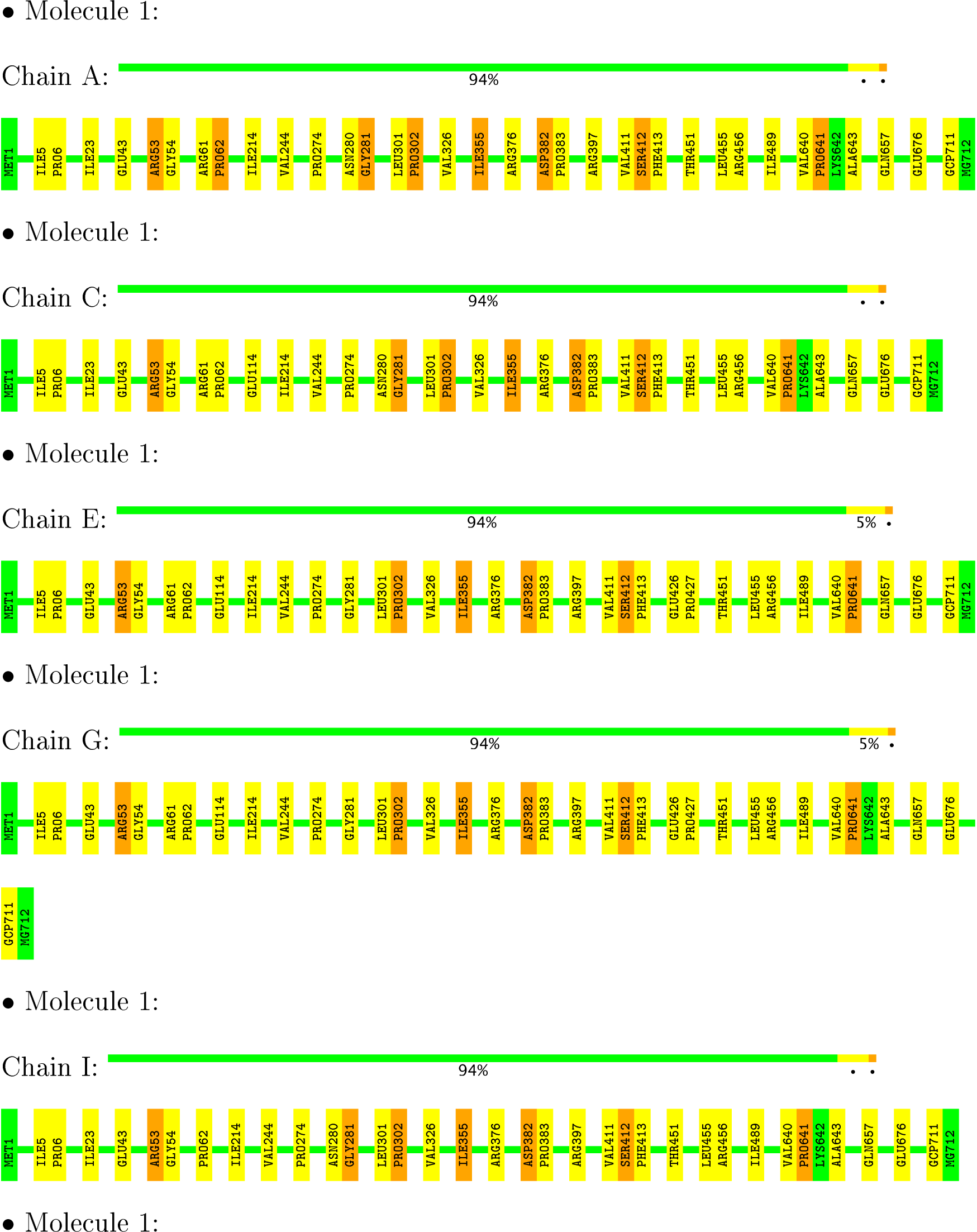
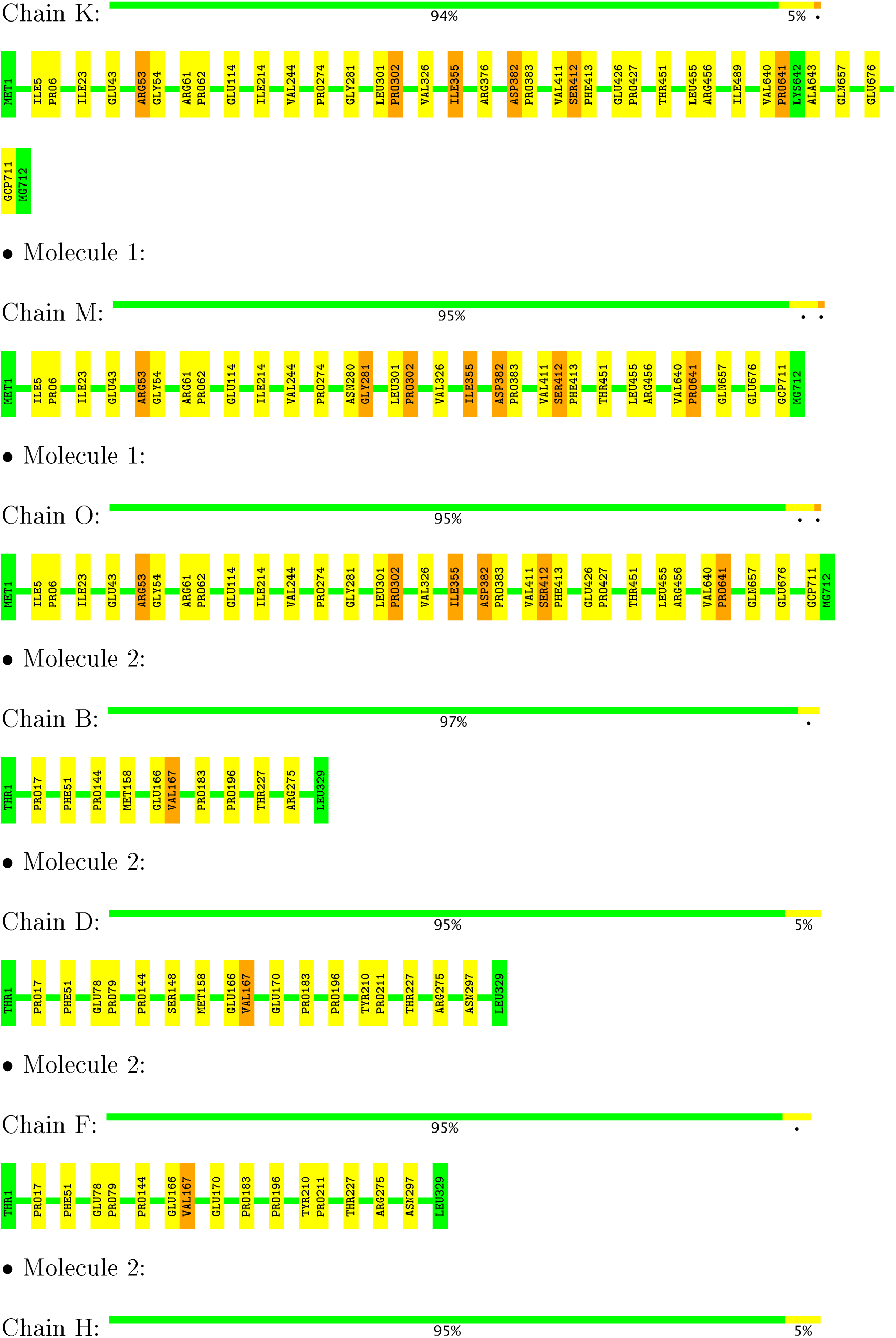
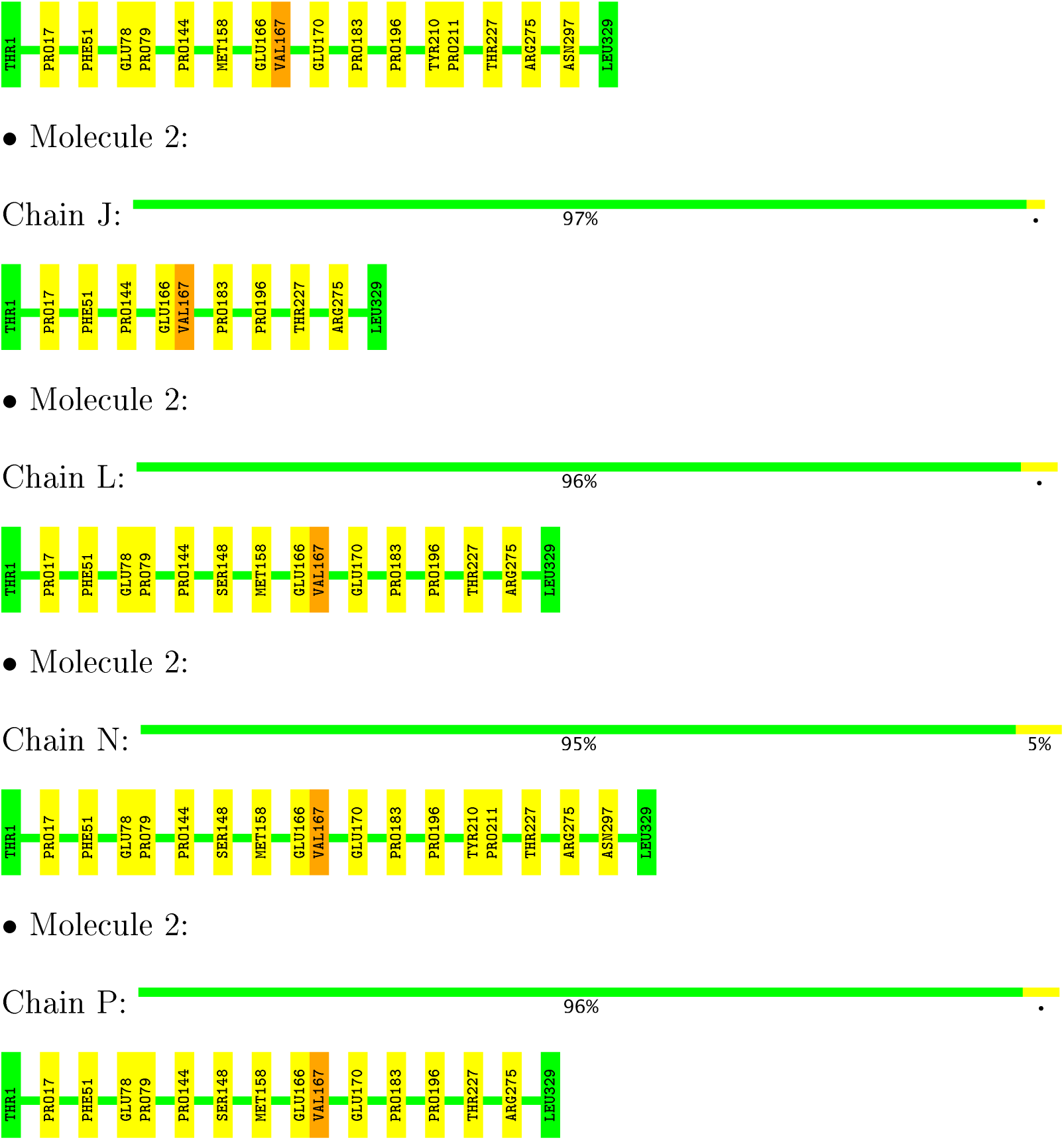

## 4 Experimental information ⓘ

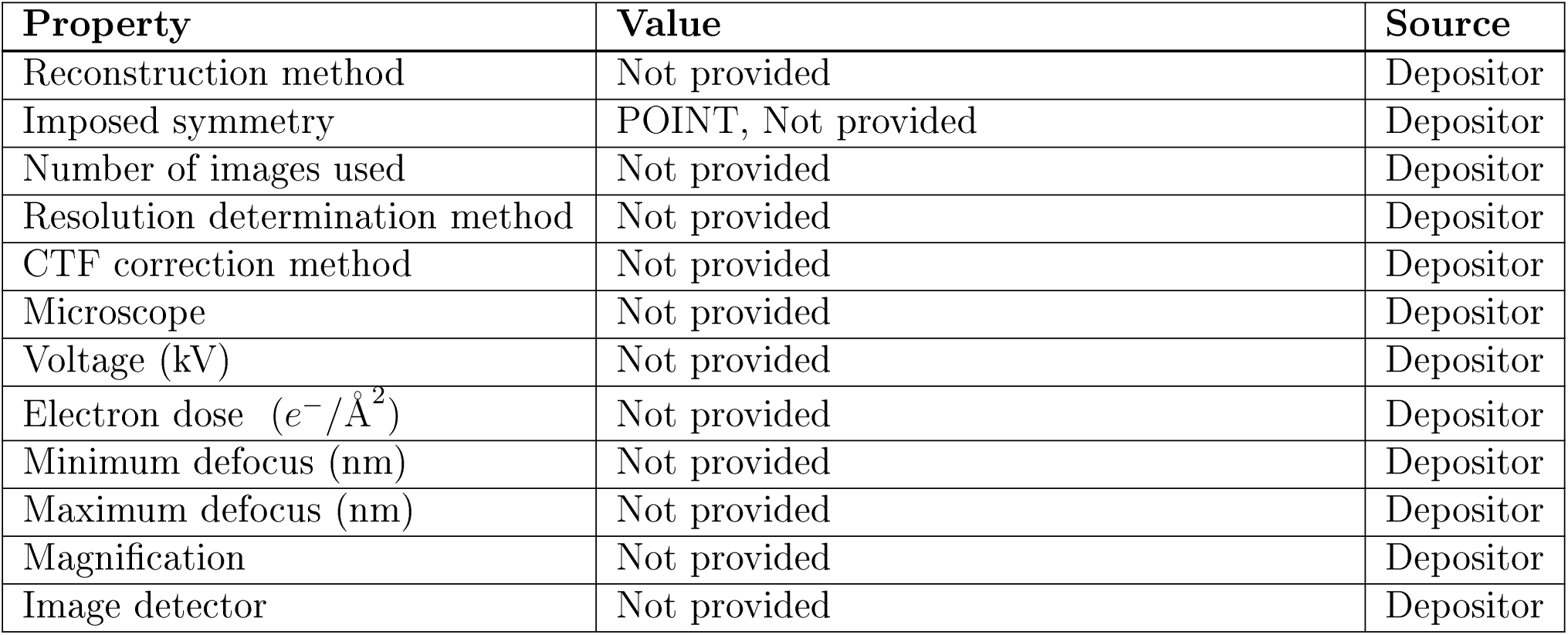

## 5 Model quality ⓘ

### 5.1 Standard geometry ⓘ

Bond lengths and bond angles in the following residue types are not validated in this section: GCP, MG

The Z score for a bond length (or angle) is the number of standard deviations the observed value is removed from the expected value. A bond length (or angle) with |*Z*| > 5 is considered an outlier worth inspection. RMSZ is the root-mean-square of all Z scores of the bond lengths (or angles).

| Mol | Chain | Bond lengths |  | Bond angles |  |
| --- | --- | --- | --- | --- | --- |
|  |  | RMSZ | #\|Z\| >2 | RMSZ | #\|Z\| >2 |
| 1 | A | 0.72 | 0/4727 | 0.51 | 0/6400 |
| 1 | C | 0.72 | 0/4727 | 0.51 | 0/6400 |
| 1 | E | 0.72 | 0/4727 | 0.51 | 0/6400 |
| 1 | G | 0.72 | 0/4727 | 0.51 | 0/6400 |
| 1 | I | 0.72 | 0/4727 | 0.51 | 0/6400 |
| 1 | K | 0.72 | 0/4727 | 0.51 | 0/6400 |
| 1 | M | 0.72 | 0/4727 | 0.51 | 0/6400 |
| 1 | O | 0.72 | 0/4727 | 0.51 | 0/6400 |
| 2 | B | 0.72 | 0/2601 | 0.49 | 0/3543 |
| 2 | D | 0.72 | 0/2601 | 0.49 | 0/3543 |
| 2 | F | 0.73 | 0/2601 | 0.49 | 0/3543 |
| 2 | H | 0.72 | 0/2601 | 0.49 | 0/3543 |
| 2 | J | 0.72 | 0/2601 | 0.49 | 0/3543 |
| 2 | L | 0.72 | 0/2601 | 0.49 | 0/3543 |
| 2 | N | 0.72 | 0/2601 | 0.49 | 0/3543 |
| 2 | P | 0.72 | 0/2601 | 0.49 | 0/3543 |
| All | All | 0.72 | 0/58624 | 0.50 | 0/79544 |

There are no bond length outliers.

There are no bond angle outliers.

There are no chirality outliers.

There are no planarity outliers.

### 5.2 Too-close contacts ⓘ

In the following table, the Non-H and H(model) columns list the number of non-hydrogen atoms and hydrogen atoms in the chain respectively. The H(added) column lists the number of hydrogen atoms added and optimized by MolProbity. The Clashes column lists the number of clashes within the asymmetric unit, whereas Symm-Clashes lists symmetry related clashes.

| Mol | Chain | Non-H | H(model) | H(added) | Clashes | Symm-Clashes |
| --- | --- | --- | --- | --- | --- | --- |
| 1 | A | 4692 | 4801 | 4813 | 27 | 0 |
| 1 | C | 4692 | 4801 | 4813 | 25 | 0 |
| 1 | E | 4692 | 4801 | 4813 | 24 | 0 |
| 1 | G | 4692 | 4801 | 4813 | 26 | 0 |
| 1 | I | 4692 | 4801 | 4813 | 25 | 0 |
| 1 | K | 4692 | 4801 | 4813 | 26 | 0 |
| 1 | M | 4692 | 4801 | 4813 | 23 | 0 |
| 1 | O | 4692 | 4801 | 4813 | 23 | 0 |
| 2 | B | 2548 | 2579 | 2581 | 5 | 0 |
| 2 | D | 2548 | 2579 | 2581 | 10 | 0 |
| 2 | F | 2548 | 2579 | 2581 | 8 | 0 |
| 2 | H | 2548 | 2579 | 2581 | 9 | 0 |
| 2 | J | 2548 | 2579 | 2581 | 5 | 0 |
| 2 | L | 2548 | 2579 | 2581 | 9 | 0 |
| 2 | N | 2548 | 2579 | 2581 | 10 | 0 |
| 2 | P | 2548 | 2579 | 2581 | 9 | 0 |
| All | All | 57920 | 59040 | 59152 | 249 | 0 |

The all-atom clashscore is defined as the number of clashes found per 1000 atoms (including hydrogen atoms). The all-atom clashscore for this structure is 2.

All (249) close contacts within the same asymmetric unit are listed below, sorted by their clash magnitude.

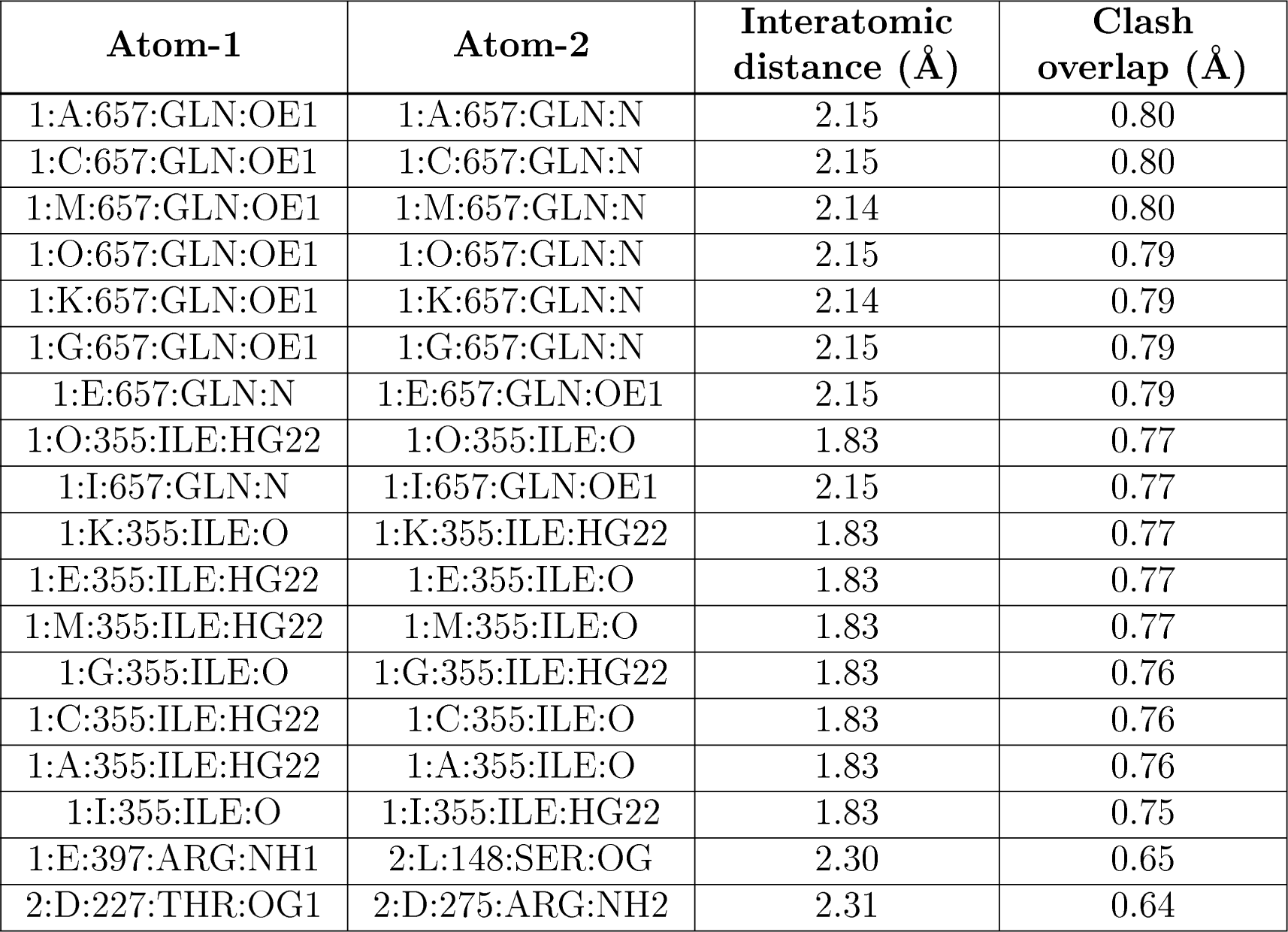
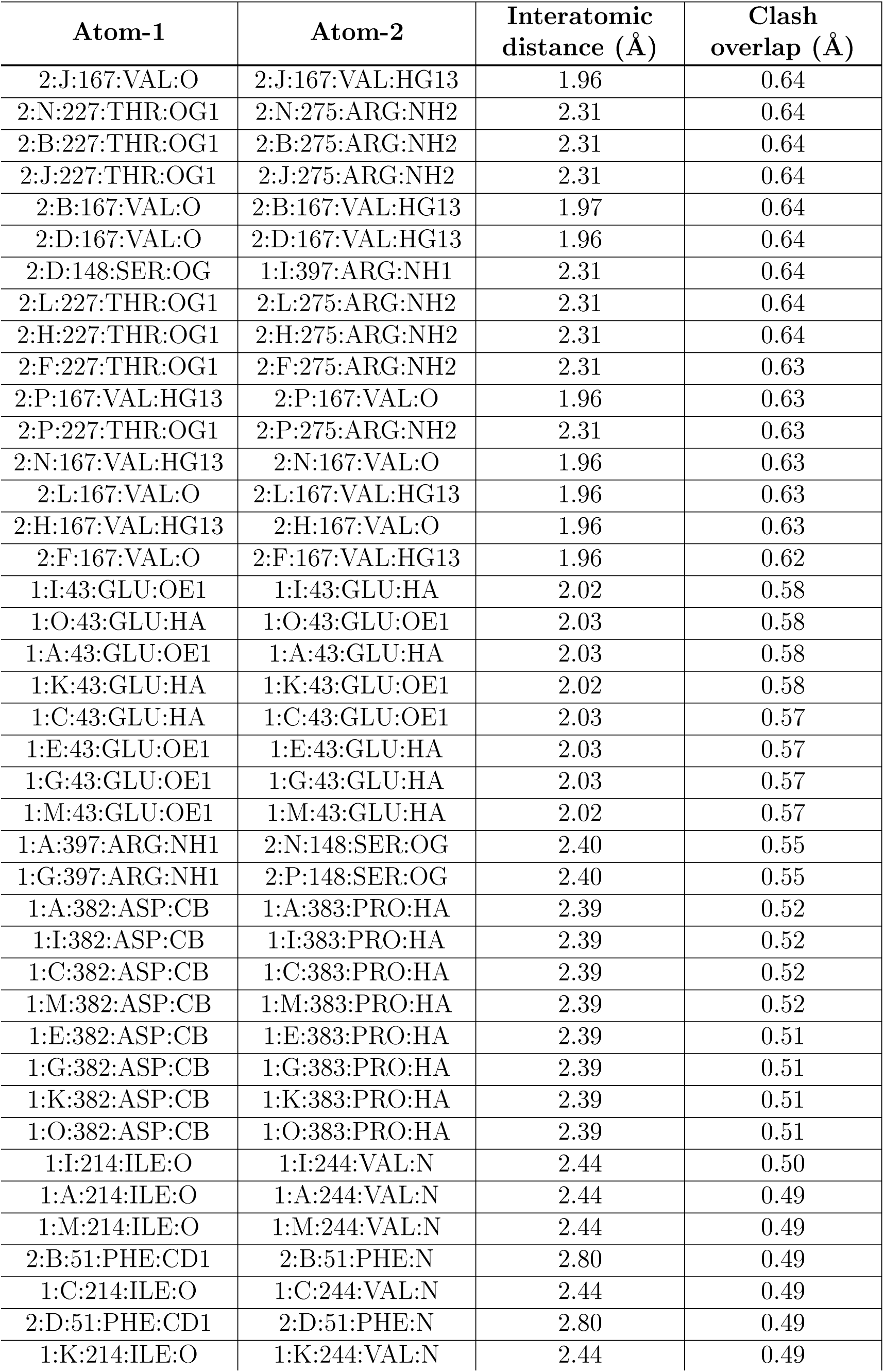
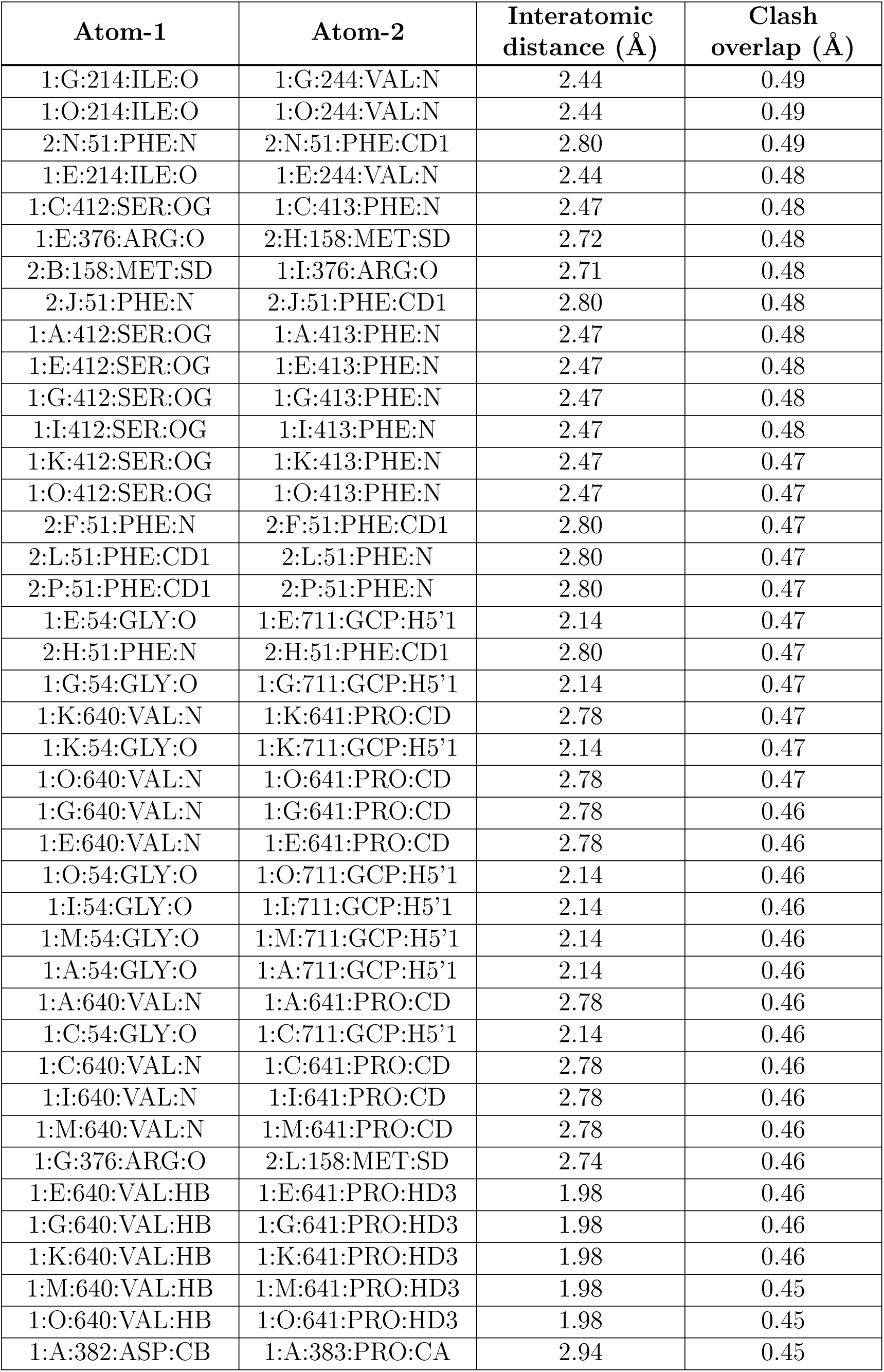
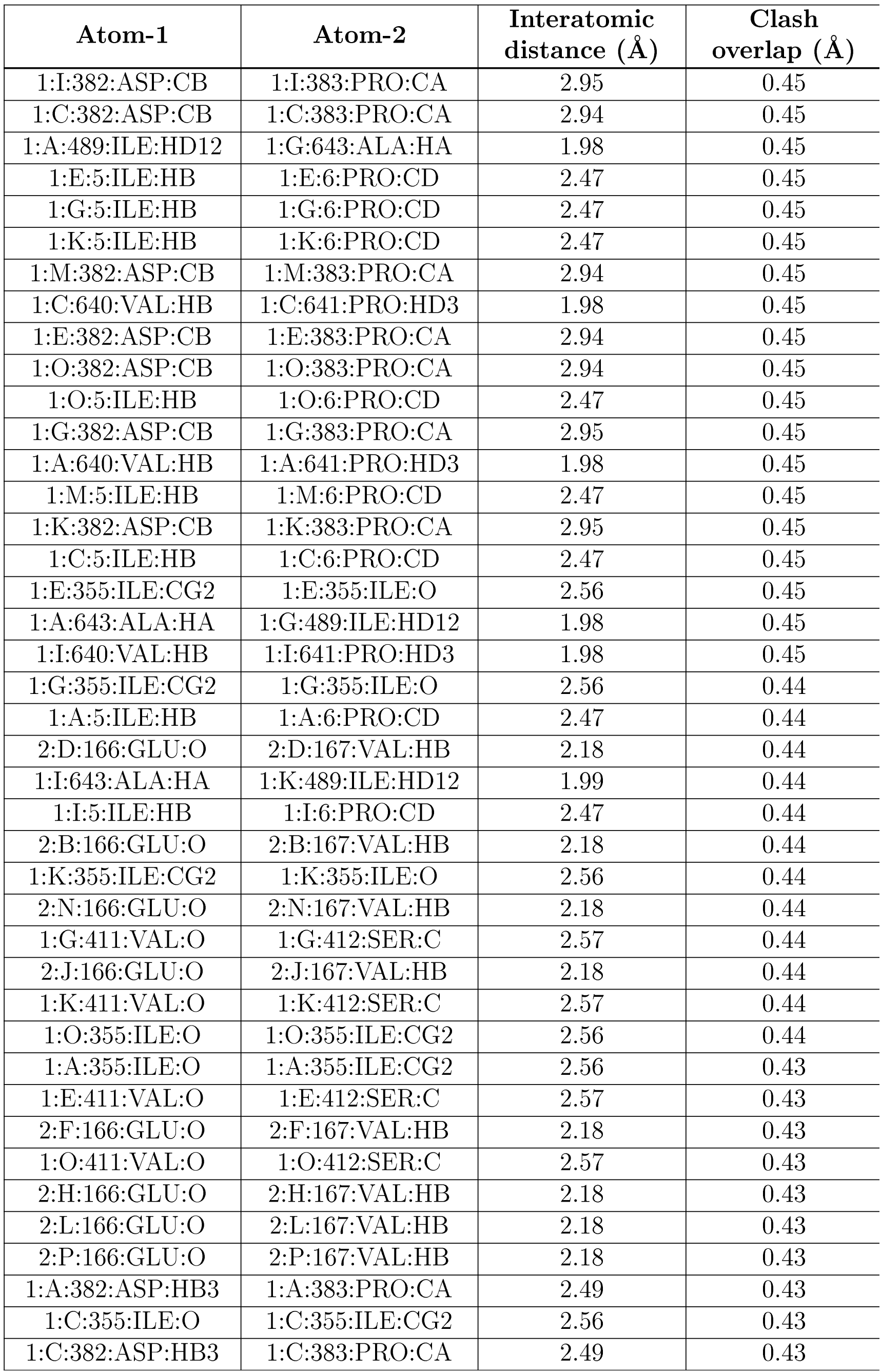
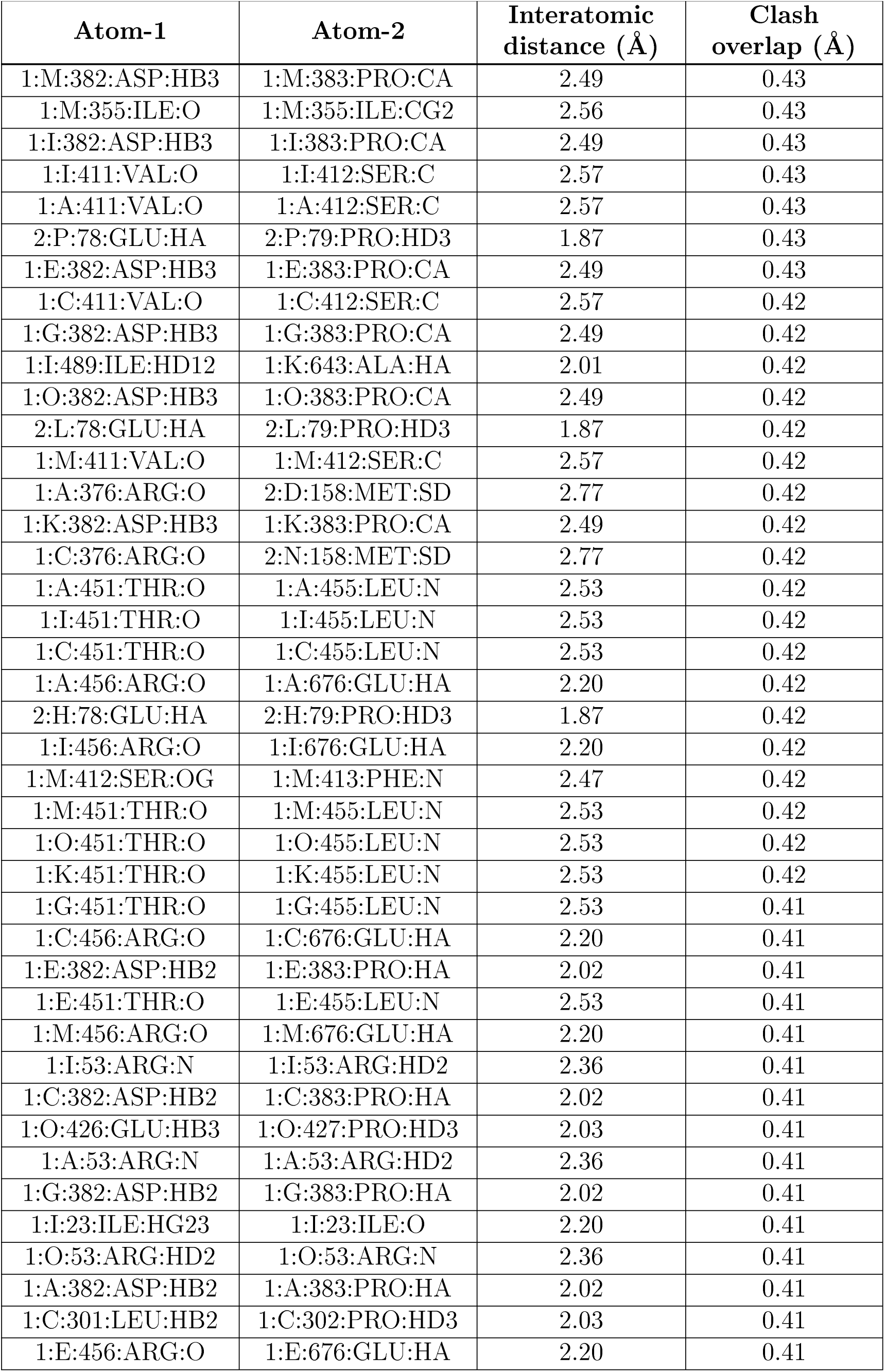
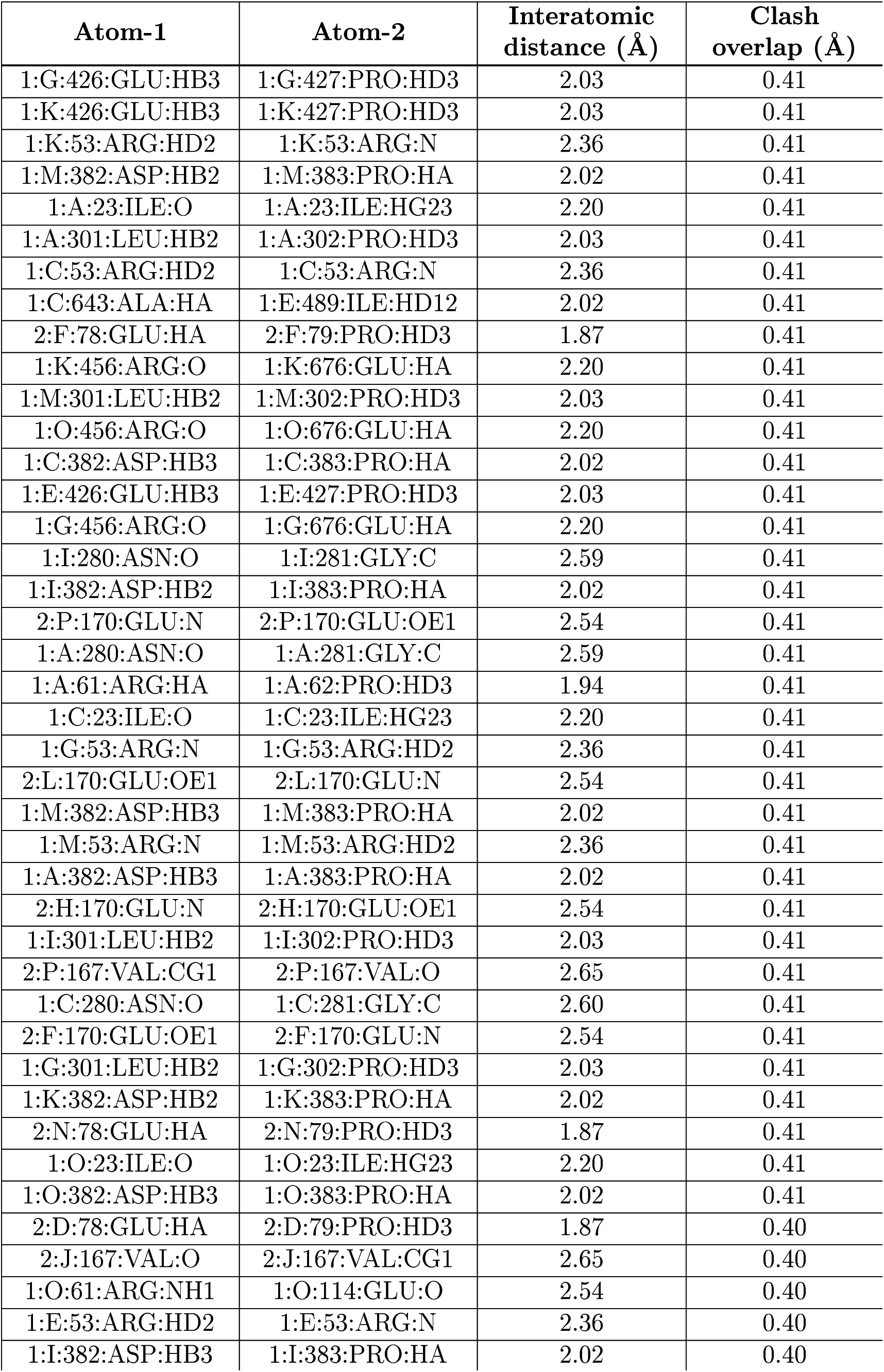
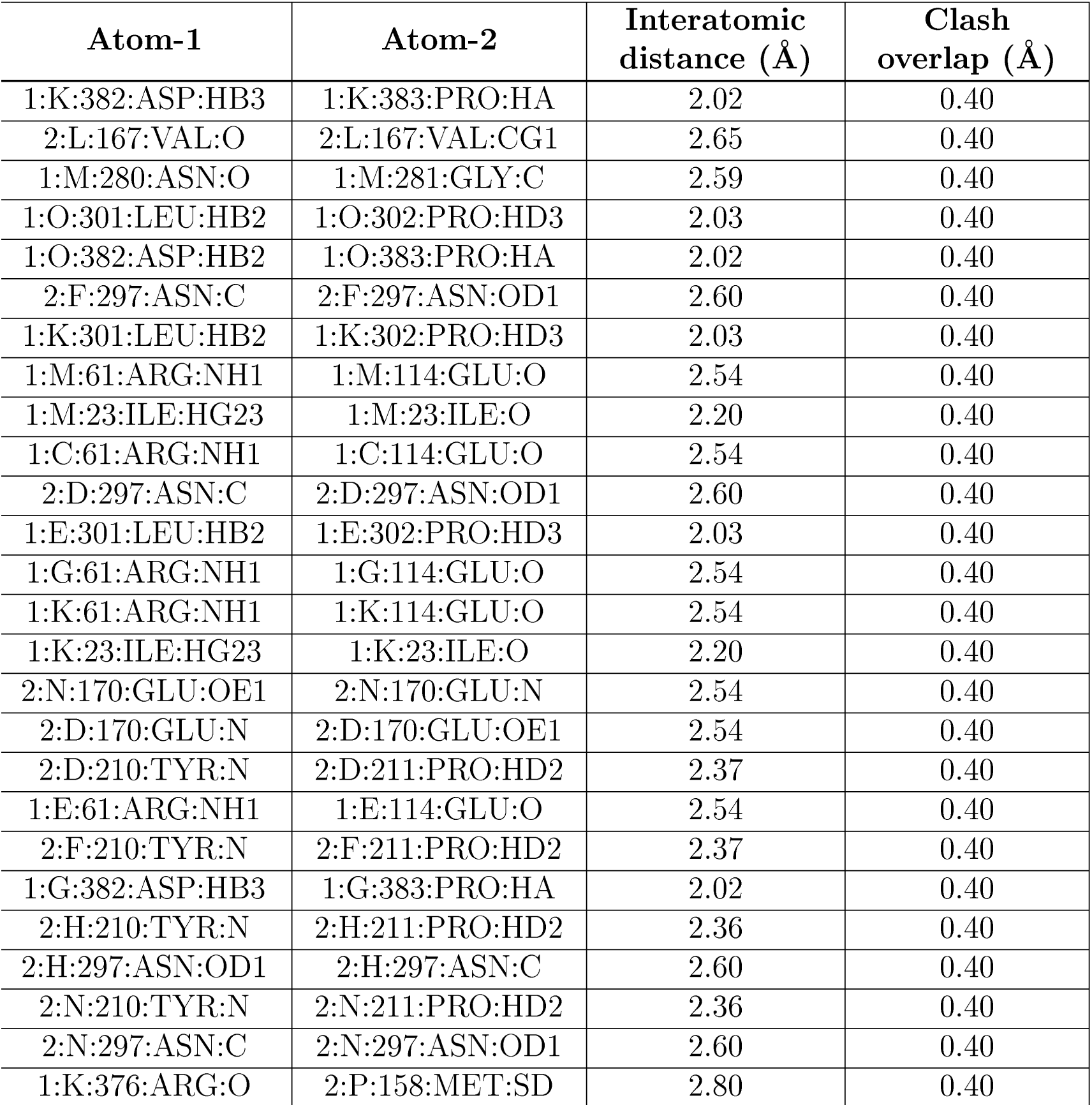

| Atom-1 | Atom-2 | Interatomic distance (Å) | Clash overlap (Å) |
| --- | --- | --- | --- |
| 1:A:657:GLN:OE1 | 1:A:657:GLN:N | 2.15 | 0.80 |
| 1:C:657:GLN:OE1 | 1:C:657:GLN:N | 2.15 | 0.80 |
| 1:M:657:GLN:OE1 | 1:M:657:GLN:N | 2.14 | 0.80 |
| 1:O:657:GLN:OE1 | 1:O:657:GLN:N | 2.15 | 0.79 |
| 1:K:657:GLN:OE1 | 1:K:657:GLN:N | 2.14 | 0.79 |
| 1:G:657:GLN:OE1 | 1:G:657:GLN:N | 2.15 | 0.79 |
| 1:E:657:GLN:N | 1:E:657:GLN:OE1 | 2.15 | 0.79 |
| 1:O:355:ILE:HG22 | 1:O:355:ILE:O | 1.83 | 0.77 |
| 1:I:657:GLN:N | 1:I:657:GLN:OE1 | 2.15 | 0.77 |
| 1:K:355:ILE:O | 1:K:355:ILE:HG22 | 1.83 | 0.77 |
| 1:E:355:ILE:HG22 | 1:E:355:ILE:O | 1.83 | 0.77 |
| 1:M:355:ILE:HG22 | 1:M:355:ILE:O | 1.83 | 0.77 |
| 1:G:355:ILE:O | 1:G:355:ILE:HG22 | 1.83 | 0.76 |
| 1:C:355:ILE:HG22 | 1:C:355:ILE:O | 1.83 | 0.76 |
| 1:A:355:ILE:HG22 | 1:A:355:ILE:O | 1.83 | 0.76 |
| 1:I:355:ILE:O | 1:I:355:ILE:HG22 | 1.83 | 0.75 |
| 1:E:397:ARG:NH1 | 2:L:148:SER:OG | 2.30 | 0.65 |
| 2:D:227:THR:OG1 | 2:D:275:ARG:NH2 | 2.31 | 0.64 |
| 2:J:167:VAL:O | 2:J:167:VAL:HG13 | 1.96 | 0.64 |
| 2:N:227:THR:OG1 | 2:N:275:ARG:NH2 | 2.31 | 0.64 |
| 2:B:227:THR:OG1 | 2:B:275:ARG:NH2 | 2.31 | 0.64 |
| 2:J:227:THR:OG1 | 2:J:275:ARG:NH2 | 2.31 | 0.64 |
| 2:B:167:VAL:O | 2:B:167:VAL:HG13 | 1.97 | 0.64 |
| 2:D:167:VAL:O | 2:D:167:VAL:HG13 | 1.96 | 0.64 |
| 2:D:148:SER:OG | 1:I:397:ARG:NH1 | 2.31 | 0.64 |
| 2:L:227:THR:OG1 | 2:L:275:ARG:NH2 | 2.31 | 0.64 |
| 2:H:227:THR:OG1 | 2:H:275:ARG:NH2 | 2.31 | 0.64 |
| 2:F:227:THR:OG1 | 2:F:275:ARG:NH2 | 2.31 | 0.63 |
| 2:P:167:VAL:HG13 | 2:P:167:VAL:O | 1.96 | 0.63 |
| 2:P:227:THR:OG1 | 2:P:275:ARG:NH2 | 2.31 | 0.63 |
| 2:N:167:VAL:HG13 | 2:N:167:VAL:O | 1.96 | 0.63 |
| 2:L:167:VAL:O | 2:L:167:VAL:HG13 | 1.96 | 0.63 |
| 2:H:167:VAL:HG13 | 2:H:167:VAL:O | 1.96 | 0.63 |
| 2:F:167:VAL:O | 2:F:167:VAL:HG13 | 1.96 | 0.62 |
| 1:I:43:GLU:OE1 | 1:I:43:GLU:HA | 2.02 | 0.58 |
| 1:O:43:GLU:HA | 1:O:43:GLU:OE1 | 2.03 | 0.58 |
| 1:A:43:GLU:OE1 | 1:A:43:GLU:HA | 2.03 | 0.58 |
| 1:K:43:GLU:HA | 1:K:43:GLU:OE1 | 2.02 | 0.58 |
| 1:C:43:GLU:HA | 1:C:43:GLU:OE1 | 2.03 | 0.57 |
| 1:E:43:GLU:OE1 | 1:E:43:GLU:HA | 2.03 | 0.57 |
| 1:G:43:GLU:OE1 | 1:G:43:GLU:HA | 2.03 | 0.57 |
| 1:M:43:GLU:OE1 | 1:M:43:GLU:HA | 2.02 | 0.57 |
| 1:A:397:ARG:NH1 | 2:N:148:SER:OG | 2.40 | 0.55 |
| 1:G:397:ARG:NH1 | 2:P:148:SER:OG | 2.40 | 0.55 |
| 1:A:382:ASP:CB | 1:A:383:PRO:HA | 2.39 | 0.52 |
| 1:I:382:ASP:CB | 1:I:383:PRO:HA | 2.39 | 0.52 |
| 1:C:382:ASP:CB | 1:C:383:PRO:HA | 2.39 | 0.52 |
| 1:M:382:ASP:CB | 1:M:383:PRO:HA | 2.39 | 0.52 |
| 1:E:382:ASP:CB | 1:E:383:PRO:HA | 2.39 | 0.51 |
| 1:G:382:ASP:CB | 1:G:383:PRO:HA | 2.39 | 0.51 |
| 1:K:382:ASP:CB | 1:K:383:PRO:HA | 2.39 | 0.51 |
| 1:O:382:ASP:CB | 1:O:383:PRO:HA | 2.39 | 0.51 |
| 1:I:214:ILE:O | 1:I:244:VAL:N | 2.44 | 0.50 |
| 1:A:214:ILE:O | 1:A:244:VAL:N | 2.44 | 0.49 |
| 1:M:214:ILE:O | 1:M:244:VAL:N | 2.44 | 0.49 |
| 2:B:51:PHE:CD1 | 2:B:51:PHE:N | 2.80 | 0.49 |
| 1:C:214:ILE:O | 1:C:244:VAL:N | 2.44 | 0.49 |
| 2:D:51:PHE:CD1 | 2:D:51:PHE:N | 2.80 | 0.49 |
| 1:K:214:ILE:O | 1:K:244:VAL:N | 2.44 | 0.49 |
| 1:G:214:ILE:O | 1:G:244:VAL:N | 2.44 | 0.49 |
| 1:O:214:ILE:O | 1:O:244:VAL:N | 2.44 | 0.49 |
| 2:N:51:PHE:N | 2:N:51:PHE:CD1 | 2.80 | 0.49 |
| 1:E:214:ILE:O | 1:E:244:VAL:N | 2.44 | 0.48 |
| 1:C:412:SER:OG | 1:C:413:PHE:N | 2.47 | 0.48 |
| 1:E:376:ARG:O | 2:H:158:MET:SD | 2.72 | 0.48 |
| 2:B:158:MET:SD | 1:I:376:ARG:O | 2.71 | 0.48 |
| 2:J:51:PHE:N | 2:J:51:PHE:CD1 | 2.80 | 0.48 |
| 1:A:412:SER:OG | 1:A:413:PHE:N | 2.47 | 0.48 |
| 1:E:412:SER:OG | 1:E:413:PHE:N | 2.47 | 0.48 |
| 1:G:412:SER:OG | 1:G:413:PHE:N | 2.47 | 0.48 |
| 1:I:412:SER:OG | 1:I:413:PHE:N | 2.47 | 0.48 |
| 1:K:412:SER:OG | 1:K:413:PHE:N | 2.47 | 0.47 |
| 1:O:412:SER:OG | 1:O:413:PHE:N | 2.47 | 0.47 |
| 2:F:51:PHE:N | 2:F:51:PHE:CD1 | 2.80 | 0.47 |
| 2:L:51:PHE:CD1 | 2:L:51:PHE:N | 2.80 | 0.47 |
| 2:P:51:PHE:CD1 | 2:P:51:PHE:N | 2.80 | 0.47 |
| 1:E:54:GLY:O | 1:E:711:GCP:H5'1 | 2.14 | 0.47 |
| 2:H:51:PHE:N | 2:H:51:PHE:CD1 | 2.80 | 0.47 |
| 1:G:54:GLY:O | 1:G:711:GCP:H5'1 | 2.14 | 0.47 |
| 1:K:640:VAL:N | 1:K:641:PRO:CD | 2.78 | 0.47 |
| 1:K:54:GLY:O | 1:K:711:GCP:H5'1 | 2.14 | 0.47 |
| 1:O:640:VAL:N | 1:O:641:PRO:CD | 2.78 | 0.47 |
| 1:G:640:VAL:N | 1:G:641:PRO:CD | 2.78 | 0.46 |
| 1:E:640:VAL:N | 1:E:641:PRO:CD | 2.78 | 0.46 |
| 1:O:54:GLY:O | 1:O:711:GCP:H5'1 | 2.14 | 0.46 |
| 1:I:54:GLY:O | 1:I:711:GCP:H5'1 | 2.14 | 0.46 |
| 1:M:54:GLY:O | 1:M:711:GCP:H5'1 | 2.14 | 0.46 |
| 1:A:54:GLY:O | 1:A:711:GCP:H5'1 | 2.14 | 0.46 |
| 1:A:640:VAL:N | 1:A:641:PRO:CD | 2.78 | 0.46 |
| 1:C:54:GLY:O | 1:C:711:GCP:H5'1 | 2.14 | 0.46 |
| 1:C:640:VAL:N | 1:C:641:PRO:CD | 2.78 | 0.46 |
| 1:I:640:VAL:N | 1:I:641:PRO:CD | 2.78 | 0.46 |
| 1:M:640:VAL:N | 1:M:641:PRO:CD | 2.78 | 0.46 |
| 1:G:376:ARG:O | 2:L:158:MET:SD | 2.74 | 0.46 |
| 1:E:640:VAL:HB | 1:E:641:PRO:HD3 | 1.98 | 0.46 |
| 1:G:640:VAL:HB | 1:G:641:PRO:HD3 | 1.98 | 0.46 |
| 1:K:640:VAL:HB | 1:K:641:PRO:HD3 | 1.98 | 0.46 |
| 1:M:640:VAL:HB | 1:M:641:PRO:HD3 | 1.98 | 0.45 |
| 1:O:640:VAL:HB | 1:O:641:PRO:HD3 | 1.98 | 0.45 |
| 1:A:382:ASP:CB | 1:A:383:PRO:CA | 2.94 | 0.45 |
| 1:I:382:ASP:CB | 1:I:383:PRO:CA | 2.95 | 0.45 |
| 1:C:382:ASP:CB | 1:C:383:PRO:CA | 2.94 | 0.45 |
| 1:A:489:ILE:HD12 | 1:G:643:ALA:HA | 1.98 | 0.45 |
| 1:E:5:ILE:HB | 1:E:6:PRO:CD | 2.47 | 0.45 |
| 1:G:5:ILE:HB | 1:G:6:PRO:CD | 2.47 | 0.45 |
| 1:K:5:ILE:HB | 1:K:6:PRO:CD | 2.47 | 0.45 |
| 1:M:382:ASP:CB | 1:M:383:PRO:CA | 2.94 | 0.45 |
| 1:C:640:VAL:HB | 1:C:641:PRO:HD3 | 1.98 | 0.45 |
| 1:E:382:ASP:CB | 1:E:383:PRO:CA | 2.94 | 0.45 |
| 1:O:382:ASP:CB | 1:O:383:PRO:CA | 2.94 | 0.45 |
| 1:O:5:ILE:HB | 1:O:6:PRO:CD | 2.47 | 0.45 |
| 1:G:382:ASP:CB | 1:G:383:PRO:CA | 2.95 | 0.45 |
| 1:A:640:VAL:HB | 1:A:641:PRO:HD3 | 1.98 | 0.45 |
| 1:M:5:ILE:HB | 1:M:6:PRO:CD | 2.47 | 0.45 |
| 1:K:382:ASP:CB | 1:K:383:PRO:CA | 2.95 | 0.45 |
| 1:C:5:ILE:HB | 1:C:6:PRO:CD | 2.47 | 0.45 |
| 1:E:355:ILE:CG2 | 1:E:355:ILE:O | 2.56 | 0.45 |
| 1:A:643:ALA:HA | 1:G:489:ILE:HD12 | 1.98 | 0.45 |
| 1:I:640:VAL:HB | 1:I:641:PRO:HD3 | 1.98 | 0.45 |
| 1:G:355:ILE:CG2 | 1:G:355:ILE:O | 2.56 | 0.44 |
| 1:A:5:ILE:HB | 1:A:6:PRO:CD | 2.47 | 0.44 |
| 2:D:166:GLU:O | 2:D:167:VAL:HB | 2.18 | 0.44 |
| 1:I:643:ALA:HA | 1:K:489:ILE:HD12 | 1.99 | 0.44 |
| 1:I:5:ILE:HB | 1:I:6:PRO:CD | 2.47 | 0.44 |
| 2:B:166:GLU:O | 2:B:167:VAL:HB | 2.18 | 0.44 |
| 1:K:355:ILE:CG2 | 1:K:355:ILE:O | 2.56 | 0.44 |
| 2:N:166:GLU:O | 2:N:167:VAL:HB | 2.18 | 0.44 |
| 1:G:411:VAL:O | 1:G:412:SER:C | 2.57 | 0.44 |
| 2:J:166:GLU:O | 2:J:167:VAL:HB | 2.18 | 0.44 |
| 1:K:411:VAL:O | 1:K:412:SER:C | 2.57 | 0.44 |
| 1:O:355:ILE:O | 1:O:355:ILE:CG2 | 2.56 | 0.44 |
| 1:A:355:ILE:O | 1:A:355:ILE:CG2 | 2.56 | 0.43 |
| 1:E:411:VAL:O | 1:E:412:SER:C | 2.57 | 0.43 |
| 2:F:166:GLU:O | 2:F:167:VAL:HB | 2.18 | 0.43 |
| 1:O:411:VAL:O | 1:O:412:SER:C | 2.57 | 0.43 |
| 2:H:166:GLU:O | 2:H:167:VAL:HB | 2.18 | 0.43 |
| 2:L:166:GLU:O | 2:L:167:VAL:HB | 2.18 | 0.43 |
| 2:P:166:GLU:O | 2:P:167:VAL:HB | 2.18 | 0.43 |
| 1:A:382:ASP:HB3 | 1:A:383:PRO:CA | 2.49 | 0.43 |
| 1:C:355:ILE:O | 1:C:355:ILE:CG2 | 2.56 | 0.43 |
| 1:C:382:ASP:HB3 | 1:C:383:PRO:CA | 2.49 | 0.43 |
| 1:M:382:ASP:HB3 | 1:M:383:PRO:CA | 2.49 | 0.43 |
| 1:M:355:ILE:O | 1:M:355:ILE:CG2 | 2.56 | 0.43 |
| 1:I:382:ASP:HB3 | 1:I:383:PRO:CA | 2.49 | 0.43 |
| 1:I:411:VAL:O | 1:I:412:SER:C | 2.57 | 0.43 |
| 1:A:411:VAL:O | 1:A:412:SER:C | 2.57 | 0.43 |
| 2:P:78:GLU:HA | 2:P:79:PRO:HD3 | 1.87 | 0.43 |
| 1:E:382:ASP:HB3 | 1:E:383:PRO:CA | 2.49 | 0.43 |
| 1:C:411:VAL:O | 1:C:412:SER:C | 2.57 | 0.42 |
| 1:G:382:ASP:HB3 | 1:G:383:PRO:CA | 2.49 | 0.42 |
| 1:I:489:ILE:HD12 | 1:K:643:ALA:HA | 2.01 | 0.42 |
| 1:O:382:ASP:HB3 | 1:O:383:PRO:CA | 2.49 | 0.42 |
| 2:L:78:GLU:HA | 2:L:79:PRO:HD3 | 1.87 | 0.42 |
| 1:M:411:VAL:O | 1:M:412:SER:C | 2.57 | 0.42 |
| 1:A:376:ARG:O | 2:D:158:MET:SD | 2.77 | 0.42 |
| 1:K:382:ASP:HB3 | 1:K:383:PRO:CA | 2.49 | 0.42 |
| 1:C:376:ARG:O | 2:N:158:MET:SD | 2.77 | 0.42 |
| 1:A:451:THR:O | 1:A:455:LEU:N | 2.53 | 0.42 |
| 1:I:451:THR:O | 1:I:455:LEU:N | 2.53 | 0.42 |
| 1:C:451:THR:O | 1:C:455:LEU:N | 2.53 | 0.42 |
| 1:A:456:ARG:O | 1:A:676:GLU:HA | 2.20 | 0.42 |
| 2:H:78:GLU:HA | 2:H:79:PRO:HD3 | 1.87 | 0.42 |
| 1:I:456:ARG:O | 1:I:676:GLU:HA | 2.20 | 0.42 |
| 1:M:412:SER:OG | 1:M:413:PHE:N | 2.47 | 0.42 |
| 1:M:451:THR:O | 1:M:455:LEU:N | 2.53 | 0.42 |
| 1:O:451:THR:O | 1:O:455:LEU:N | 2.53 | 0.42 |
| 1:K:451:THR:O | 1:K:455:LEU:N | 2.53 | 0.42 |
| 1:G:451:THR:O | 1:G:455:LEU:N | 2.53 | 0.41 |
| 1:C:456:ARG:O | 1:C:676:GLU:HA | 2.20 | 0.41 |
| 1:E:382:ASP:HB2 | 1:E:383:PRO:HA | 2.02 | 0.41 |
| 1:E:451:THR:O | 1:E:455:LEU:N | 2.53 | 0.41 |
| 1:M:456:ARG:O | 1:M:676:GLU:HA | 2.20 | 0.41 |
| 1:I:53:ARG:N | 1:I:53:ARG:HD2 | 2.36 | 0.41 |
| 1:C:382:ASP:HB2 | 1:C:383:PRO:HA | 2.02 | 0.41 |
| 1:O:426:GLU:HB3 | 1:O:427:PRO:HD3 | 2.03 | 0.41 |
| 1:A:53:ARG:N | 1:A:53:ARG:HD2 | 2.36 | 0.41 |
| 1:G:382:ASP:HB2 | 1:G:383:PRO:HA | 2.02 | 0.41 |
| 1:I:23:ILE:HG23 | 1:I:23:ILE:O | 2.20 | 0.41 |
| 1:O:53:ARG:HD2 | 1:O:53:ARG:N | 2.36 | 0.41 |
| 1:A:382:ASP:HB2 | 1:A:383:PRO:HA | 2.02 | 0.41 |
| 1:C:301:LEU:HB2 | 1:C:302:PRO:HD3 | 2.03 | 0.41 |
| 1:E:456:ARG:O | 1:E:676:GLU:HA | 2.20 | 0.41 |
| 1:G:426:GLU:HB3 | 1:G:427:PRO:HD3 | 2.03 | 0.41 |
| 1:K:426:GLU:HB3 | 1:K:427:PRO:HD3 | 2.03 | 0.41 |
| 1:K:53:ARG:HD2 | 1:K:53:ARG:N | 2.36 | 0.41 |
| 1:M:382:ASP:HB2 | 1:M:383:PRO:HA | 2.02 | 0.41 |
| 1:A:23:ILE:O | 1:A:23:ILE:HG23 | 2.20 | 0.41 |
| 1:A:301:LEU:HB2 | 1:A:302:PRO:HD3 | 2.03 | 0.41 |
| 1:C:53:ARG:HD2 | 1:C:53:ARG:N | 2.36 | 0.41 |
| 1:C:643:ALA:HA | 1:E:489:ILE:HD12 | 2.02 | 0.41 |
| 2:F:78:GLU:HA | 2:F:79:PRO:HD3 | 1.87 | 0.41 |
| 1:K:456:ARG:O | 1:K:676:GLU:HA | 2.20 | 0.41 |
| 1:M:301:LEU:HB2 | 1:M:302:PRO:HD3 | 2.03 | 0.41 |
| 1:O:456:ARG:O | 1:O:676:GLU:HA | 2.20 | 0.41 |
| 1:C:382:ASP:HB3 | 1:C:383:PRO:HA | 2.02 | 0.41 |
| 1:E:426:GLU:HB3 | 1:E:427:PRO:HD3 | 2.03 | 0.41 |
| 1:G:456:ARG:O | 1:G:676:GLU:HA | 2.20 | 0.41 |
| 1:I:280:ASN:O | 1:I:281:GLY:C | 2.59 | 0.41 |
| 1:I:382:ASP:HB2 | 1:I:383:PRO:HA | 2.02 | 0.41 |
| 2:P:170:GLU:N | 2:P:170:GLU:OE1 | 2.54 | 0.41 |
| 1:A:280:ASN:O | 1:A:281:GLY:C | 2.59 | 0.41 |
| 1:A:61:ARG:HA | 1:A:62:PRO:HD3 | 1.94 | 0.41 |
| 1:C:23:ILE:O | 1:C:23:ILE:HG23 | 2.20 | 0.41 |
| 1:G:53:ARG:N | 1:G:53:ARG:HD2 | 2.36 | 0.41 |
| 2:L:170:GLU:OE1 | 2:L:170:GLU:N | 2.54 | 0.41 |
| 1:M:382:ASP:HB3 | 1:M:383:PRO:HA | 2.02 | 0.41 |
| 1:M:53:ARG:N | 1:M:53:ARG:HD2 | 2.36 | 0.41 |
| 1:A:382:ASP:HB3 | 1:A:383:PRO:HA | 2.02 | 0.41 |
| 2:H:170:GLU:N | 2:H:170:GLU:OE1 | 2.54 | 0.41 |
| 1:I:301:LEU:HB2 | 1:I:302:PRO:HD3 | 2.03 | 0.41 |
| 2:P:167:VAL:CG1 | 2:P:167:VAL:O | 2.65 | 0.41 |
| 1:C:280:ASN:O | 1:C:281:GLY:C | 2.60 | 0.41 |
| 2:F:170:GLU:OE1 | 2:F:170:GLU:N | 2.54 | 0.41 |
| 1:G:301:LEU:HB2 | 1:G:302:PRO:HD3 | 2.03 | 0.41 |
| 1:K:382:ASP:HB2 | 1:K:383:PRO:HA | 2.02 | 0.41 |
| 2:N:78:GLU:HA | 2:N:79:PRO:HD3 | 1.87 | 0.41 |
| 1:O:23:ILE:O | 1:O:23:ILE:HG23 | 2.20 | 0.41 |
| 1:O:382:ASP:HB3 | 1:O:383:PRO:HA | 2.02 | 0.41 |
| 2:D:78:GLU:HA | 2:D:79:PRO:HD3 | 1.87 | 0.40 |
| 2:J:167:VAL:O | 2:J:167:VAL:CG1 | 2.65 | 0.40 |
| 1:O:61:ARG:NH1 | 1:O:114:GLU:O | 2.54 | 0.40 |
| 1:E:53:ARG:HD2 | 1:E:53:ARG:N | 2.36 | 0.40 |
| 1:I:382:ASP:HB3 | 1:I:383:PRO:HA | 2.02 | 0.40 |
| 1:K:382:ASP:HB3 | 1:K:383:PRO:HA | 2.02 | 0.40 |
| 2:L:167:VAL:O | 2:L:167:VAL:CG1 | 2.65 | 0.40 |
| 1:M:280:ASN:O | 1:M:281:GLY:C | 2.59 | 0.40 |
| 1:O:301:LEU:HB2 | 1:O:302:PRO:HD3 | 2.03 | 0.40 |
| 1:O:382:ASP:HB2 | 1:O:383:PRO:HA | 2.02 | 0.40 |
| 2:F:297:ASN:C | 2:F:297:ASN:OD1 | 2.60 | 0.40 |
| 1:K:301:LEU:HB2 | 1:K:302:PRO:HD3 | 2.03 | 0.40 |
| 1:M:61:ARG:NH1 | 1:M:114:GLU:O | 2.54 | 0.40 |
| 1:M:23:ILE:HG23 | 1:M:23:ILE:O | 2.20 | 0.40 |
| 1:C:61:ARG:NH1 | 1:C:114:GLU:O | 2.54 | 0.40 |
| 2:D:297:ASN:C | 2:D:297:ASN:OD1 | 2.60 | 0.40 |
| 1:E:301:LEU:HB2 | 1:E:302:PRO:HD3 | 2.03 | 0.40 |
| 1:G:61:ARG:NH1 | 1:G:114:GLU:O | 2.54 | 0.40 |
| 1:K:61:ARG:NH1 | 1:K:114:GLU:O | 2.54 | 0.40 |
| 1:K:23:ILE:HG23 | 1:K:23:ILE:O | 2.20 | 0.40 |
| 2:N:170:GLU:OE1 | 2:N:170:GLU:N | 2.54 | 0.40 |
| 2:D:170:GLU:N | 2:D:170:GLU:OE1 | 2.54 | 0.40 |
| 2:D:210:TYR:N | 2:D:211:PRO:HD2 | 2.37 | 0.40 |
| 1:E:61:ARG:NH1 | 1:E:114:GLU:O | 2.54 | 0.40 |
| 2:F:210:TYR:N | 2:F:211:PRO:HD2 | 2.37 | 0.40 |
| 1:G:382:ASP:HB3 | 1:G:383:PRO:HA | 2.02 | 0.40 |
| 2:H:210:TYR:N | 2:H:211:PRO:HD2 | 2.36 | 0.40 |
| 2:H:297:ASN:OD1 | 2:H:297:ASN:C | 2.60 | 0.40 |
| 2:N:210:TYR:N | 2:N:211:PRO:HD2 | 2.36 | 0.40 |
| 2:N:297:ASN:C | 2:N:297:ASN:OD1 | 2.60 | 0.40 |
| 1:K:376:ARG:O | 2:P:158:MET:SD | 2.80 | 0.40 |

There are no symmetry-related clashes.

### 5.3 Torsion angles ⓘ

#### 5.3.1 Protein backbone ⓘ

In the following table, the Percentiles column shows the percent Ramachandran outliers of the chain as a percentile score with respect to all PDB entries followed by that with respect to all EM entries.

The Analysed column shows the number of residues for which the backbone conformation was analysed, and the total number of residues.

| Mol | Chain | Analysed | Favoured | Allowed | Outliers | Percentiles |  |
| --- | --- | --- | --- | --- | --- | --- | --- |
| 1 | A | 583/591 (99%) | 550 (94%) | 25 (4%) | 8 (1%) | 13 | 13 |
| 1 | C | 583/591 (99%) | 550 (94%) | 25 (4%) | 8 (1%) | 13 | 13 |
| 1 | E | 583/591 (99%) | 550 (94%) | 25 (4%) | 8 (1%) | 13 | 13 |
| 1 | G | 583/591 (99%) | 550 (94%) | 25 (4%) | 8 (1%) | 13 | 13 |
| 1 | I | 583/591 (99%) | 550 (94%) | 25 (4%) | 8 (1%) | 13 | 13 |
| 1 | K | 583/591 (99%) | 550 (94%) | 25 (4%) | 8 (1%) | 13 | 13 |
| 1 | M | 583/591 (99%) | 550 (94%) | 25 (4%) | 8 (1%) | 13 | 13 |
| 1 | O | 583/591 (99%) | 550 (94%) | 25 (4%) | 8 (1%) | 13 | 13 |
| 2 | B | 327/329 (99%) | 302 (92%) | 21 (6%) | 4 (1%) | 15 | 15 |
| 2 | D | 327/329 (99%) | 302 (92%) | 21 (6%) | 4 (1%) | 15 | 15 |
| 2 | F | 327/329 (99%) | 302 (92%) | 21 (6%) | 4 (1%) | 15 | 15 |
| 2 | H | 327/329 (99%) | 302 (92%) | 21 (6%) | 4 (1%) | 15 | 15 |
| 2 | J | 327/329 (99%) | 302 (92%) | 21 (6%) | 4 (1%) | 15 | 15 |
| 2 | L | 327/329 (99%) | 302 (92%) | 21 (6%) | 4 (1%) | 15 | 15 |
| 2 | N | 327/329 (99%) | 302 (92%) | 21 (6%) | 4 (1%) | 15 | 15 |
| 2 | P | 327/329 (99%) | 302 (92%) | 21 (6%) | 4 (1%) | 15 | 15 |
| All | All | 7280/7360 (99%) | 6816 (94%) | 368 (5%) | 96 (1%) | 19 | 14 |

All (96) Ramaehandran outliers are listed below:

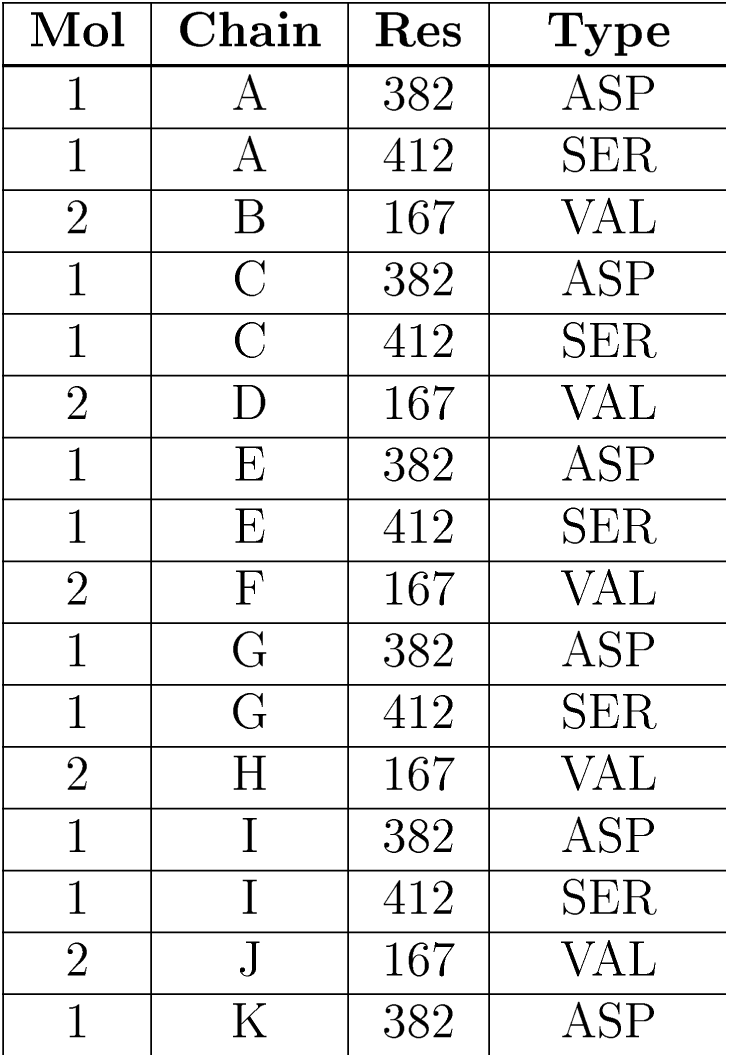
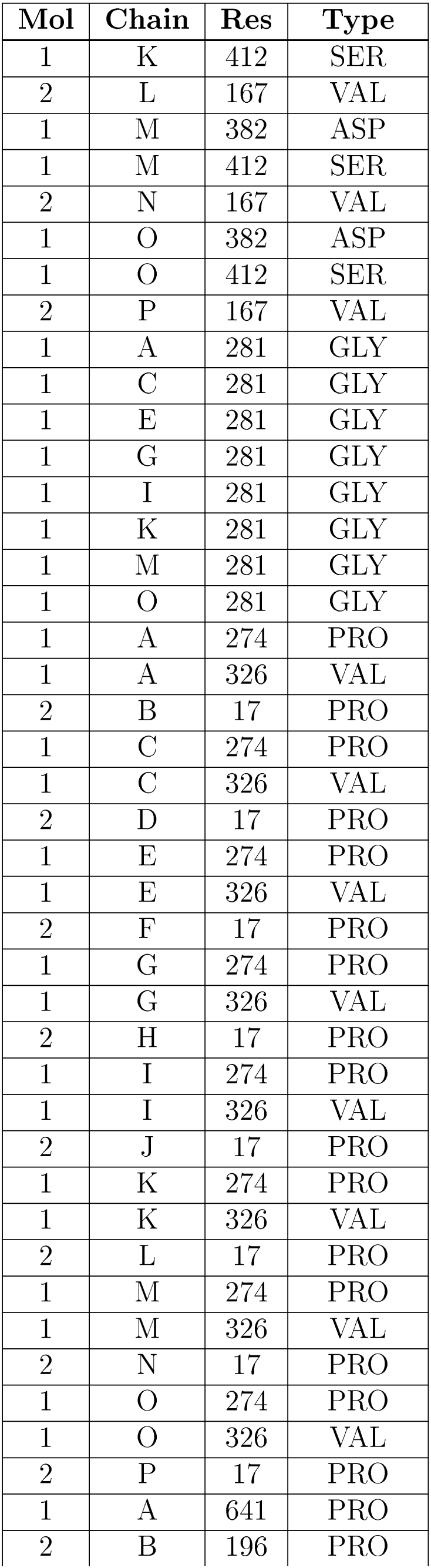
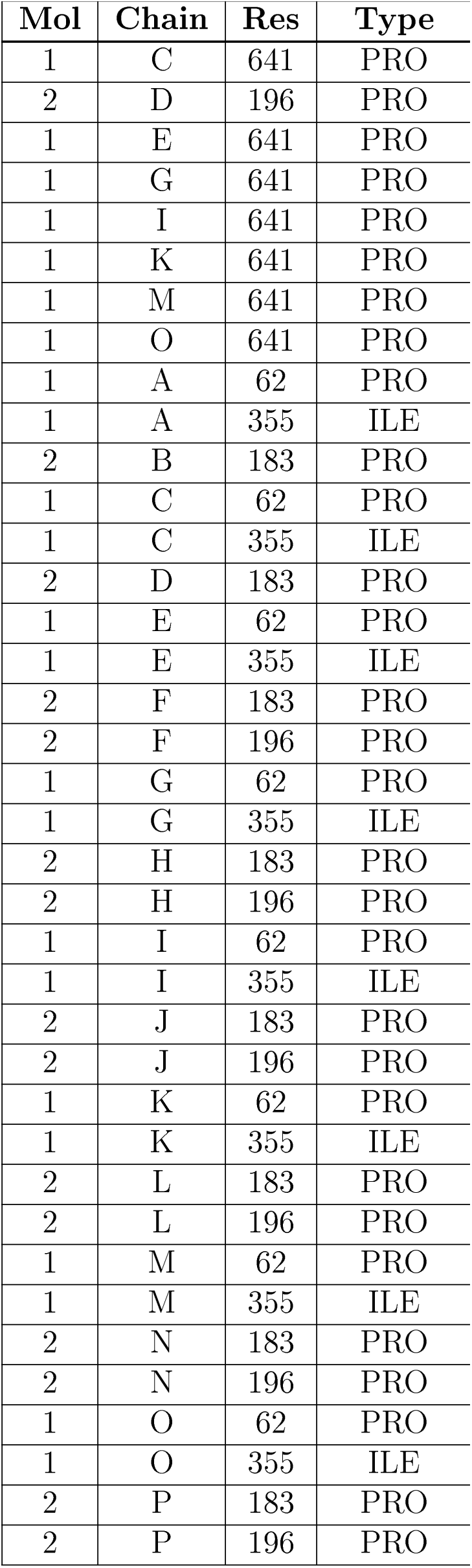

All (96) Ramachandran outliers are listed below:
| Mol | Chain | Res | Type |
| --- | --- | --- | --- |
| 1 | A | 382 | ASP |
| 1 | A | 412 | SER |
| 2 | B | 167 | VAL |
| 1 | C | 382 | ASP |
| 1 | C | 412 | SER |
| 2 | D | 167 | VAL |
| 1 | E | 382 | ASP |
| 1 | E | 412 | SER |
| 2 | F | 167 | VAL |
| 1 | G | 382 | ASP |
| 1 | G | 412 | SER |
| 2 | H | 167 | VAL |
| 1 | I | 382 | ASP |
| 1 | I | 412 | SER |
| 2 | J | 167 | VAL |
| 1 | K | 382 | ASP |
| 1 | K | 412 | SER |
| 2 | L | 167 | VAL |
| 1 | M | 382 | ASP |
| 1 | M | 412 | SER |
| 2 | N | 167 | VAL |
| 1 | O | 382 | ASP |
| 1 | O | 412 | SER |
| 2 | P | 167 | VAL |
| 1 | A | 281 | GLY |
| 1 | C | 281 | GLY |
| 1 | E | 281 | GLY |
| 1 | G | 281 | GLY |
| 1 | I | 281 | GLY |
| 1 | K | 281 | GLY |
| 1 | M | 281 | GLY |
| 1 | O | 281 | GLY |
| 1 | A | 274 | PRO |
| 1 | A | 326 | VAL |
| 2 | B | 17 | PRO |
| 1 | C | 274 | PRO |
| 1 | C | 326 | VAL |
| 2 | D | 17 | PRO |
| 1 | E | 274 | PRO |
| 1 | E | 326 | VAL |
| 2 | F | 17 | PRO |
| 1 | G | 274 | PRO |
| 1 | G | 326 | VAL |
| 2 | H | 17 | PRO |
| 1 | I | 274 | PRO |
| 1 | I | 326 | VAL |
| 2 | J | 17 | PRO |
| 1 | K | 274 | PRO |
| 1 | K | 326 | VAL |
| 2 | L | 17 | PRO |
| 1 | M | 274 | PRO |
| 1 | M | 326 | VAL |
| 2 | N | 17 | PRO |
| 1 | O | 274 | PRO |
| 1 | O | 326 | VAL |
| 2 | P | 17 | PRO |
| 1 | A | 641 | PRO |
| 2 | B | 196 | PRO |
| 1 | C | 641 | PRO |
| 2 | D | 196 | PRO |
| 1 | E | 641 | PRO |
| 1 | G | 641 | PRO |
| 1 | I | 641 | PRO |
| 1 | K | 641 | PRO |
| 1 | M | 641 | PRO |
| 1 | O | 641 | PRO |
| 1 | A | 62 | PRO |
| 1 | A | 355 | ILE |
| 2 | B | 183 | PRO |
| 1 | C | 62 | PRO |
| 1 | C | 355 | ILE |
| 2 | D | 183 | PRO |
| 1 | E | 62 | PRO |
| 1 | E | 355 | ILE |
| 2 | F | 183 | PRO |
| 2 | F | 196 | PRO |
| 1 | G | 62 | PRO |
| 1 | G | 355 | ILE |
| 2 | H | 183 | PRO |
| 2 | H | 196 | PRO |
| 1 | I | 62 | PRO |
| 1 | I | 355 | ILE |
| 2 | J | 183 | PRO |
| 2 | J | 196 | PRO |
| 1 | K | 62 | PRO |
| 1 | K | 355 | ILE |
| 2 | L | 183 | PRO |
| 2 | L | 196 | PRO |
| 1 | M | 62 | PRO |
| 1 | M | 355 | ILE |
| 2 | N | 183 | PRO |
| 2 | N | 196 | PRO |
| 1 | O | 62 | PRO |
| 1 | O | 355 | ILE |
| 2 | P | 183 | PRO |
| 2 | P | 196 | PRO |

#### 5.3.2 Protein sidechains ⓘ

In the following table, the Percentiles column shows the percent sidechain outliers of the chain as a percentile score with respect to all PDB entries followed by that with respect to all EM entries.

The Analysed column shows the number of residues for which the sidechain conformation was analysed, and the total number of residues.

| Mol | Chain | Analysed | Rotameric | Outliers | Percentiles |  |
| --- | --- | --- | --- | --- | --- | --- |
| 1 | A | 527/527 (100%) | 525 (100%) | 2 (0%) | 93 | 93 |
| 1 | C | 527/527 (100%) | 525 (100%) | 2 (0%) | 93 | 93 |
| 1 | E | 527/527 (100%) | 525 (100%) | 2 (0%) | 93 | 93 |
| 1 | G | 527/527 (100%) | 525 (100%) | 2 (0%) | 93 | 93 |
| 1 | I | 527/527 (100%) | 525 (100%) | 2 (0%) | 93 | 93 |
| 1 | K | 527/527 (100%) | 525 (100%) | 2 (0%) | 93 | 93 |
| 1 | M | 527/527 (100%) | 525 (100%) | 2 (0%) | 93 | 93 |
| 1 | O | 527/527 (100%) | 525 (100%) | 2 (0%) | 93 | 93 |
| 2 | B | 268/268 (100%) | 267 (100%) | 1 (0%) | 93 | 93 |
| 2 | D | 268/268 (100%) | 267 (100%) | 1 (0%) | 93 | 93 |
| 2 | F | 268/268 (100%) | 267 (100%) | 1 (0%) | 93 | 93 |
| 2 | H | 268/268 (100%) | 267 (100%) | 1 (0%) | 93 | 93 |
| 2 | J | 268/268 (100%) | 267 (100%) | 1 (0%) | 93 | 93 |
| 2 | L | 268/268 (100%) | 267 (100%) | 1 (0%) | 93 | 93 |
| 2 | N | 268/268 (100%) | 267 (100%) | 1 (0%) | 93 | 93 |
| 2 | P | 268/268 (100%) | 267 (100%) | 1 (0%) | 93 | 93 |
| All | All | 6360/6360 (100%) | 6336 (100%) | 24 (0%) | 93 | 93 |

All (24) residues with a non-rotamerie sideehain are listed below:

All (24) residues with a non-rotameric sidechain are listed below:
| Mol | Chain | Res | Type |
| --- | --- | --- | --- |
| 1 | A | 53 | ARG |
| 1 | A | 302 | PRO |
| 2 | B | 144 | PRO |
| 1 | C | 53 | ARG |
| 1 | C | 302 | PRO |
| 2 | D | 144 | PRO |
| 1 | E | 53 | ARG |
| 1 | E | 302 | PRO |

| Mol | Chain | Res | Type |
| --- | --- | --- | --- |
| 2 | F | 144 | PRO |
| 1 | G | 53 | ARG |
| 1 | G | 302 | PRO |
| 2 | H | 144 | PRO |
| 1 | I | 53 | ARG |
| 1 | I | 302 | PRO |
| 2 | J | 144 | PRO |
| 1 | K | 53 | ARG |
| 1 | K | 302 | PRO |
| 2 | L | 144 | PRO |
| 1 | M | 53 | ARG |
| 1 | M | 302 | PRO |
| 2 | N | 144 | PRO |
| 1 | O | 53 | ARG |
| 1 | O | 302 | PRO |
| 2 | P | 144 | PRO |

Some sidechains can be flipped to improve hydrogen bonding and reduce clashes. There are no such sidechains identified.

#### 5.3.3 RNA ⓘ

There are no RNA molecules in this entry.

### 5.4 Non-standard residues in protein, DNA, RNA chains ⓘ

There are no non-standard protein/DNA/RNA residues in this entry.

### 5.5 Carbohydrates ⓘ

There are no carbohydrates in this entry.

### 5.6 Ligand geometry ⓘ

There are no ligands in this entry.

### 5.7 Other polymers ⓘ

There are no such residues in this entry.

### 5.8 Polymer linkage issues ⓘ

The following chains have linkage breaks:

| Mol | Chain | Number of breaks |
| --- | --- | --- |
| 1 | G | 2 |
| 1 | K | 2 |
| 1 | E | 2 |
| 1 | I | 2 |
| 1 | C | 2 |
| 1 | A | 2 |
| 1 | O | 2 |
| 1 | M | 2 |

All chain breaks are listed below:

| Model | Chain | Residue-1 | Atom-1 | Residue-2 | Atom-2 | Distance (Å) |
| --- | --- | --- | --- | --- | --- | --- |
| 1 | A | 503:ASP | C | 612:ALA | N | 20.73 |
| 1 | C | 503:ASP | C | 612:ALA | N | 20.73 |
| 1 | E | 503:ASP | C | 612:ALA | N | 20.73 |
| 1 | G | 503:ASP | C | 612:ALA | N | 20.73 |
| 1 | I | 503:ASP | C | 612:ALA | N | 20.73 |
| 1 | K | 503:ASP | C | 612:ALA | N | 20.73 |
| 1 | M | 503:ASP | C | 612:ALA | N | 20.73 |
| 1 | O | 503:ASP | C | 612:ALA | N | 20.73 |
| 1 | A | 73:GLU | C | 87:ALA | N | 9.30 |
| 1 | C | 73:GLU | C | 87:ALA | N | 9.30 |
| 1 | E | 73:GLU | C | 87:ALA | N | 9.30 |
| 1 | G | 73:GLU | C | 87:ALA | N | 9.30 |
| 1 | I | 73:GLU | C | 87:ALA | N | 9.30 |
| 1 | K | 73:GLU | C | 87:ALA | N | 9.30 |
| 1 | M | 73:GLU | C | 87:ALA | N | 9.30 |
| 1 | O | 73:GLU | C | 87:ALA | N | 9.30 |

